# Quantitative molecular cartography of emergency myelopoiesis reveals conserved modules of hematopoietic activation

**DOI:** 10.1101/2025.05.28.656712

**Authors:** James W. Swann, Jun Hou Fung, Ziwei Chen, Oakley C. Olson, Amélie Collins, Tenzin Lhakhang, Melissa A. Proven, Ruiyuan Zhang, Raul Rabadan, Emmanuelle Passegué

## Abstract

Hematopoietic stem and progenitor cells (HSPC) respond to infections, inflammation, and regenerative challenges using a collection of cellular and molecular mechanisms termed emergency myelopoiesis (EM) pathways. However, it remains unclear how various EM inducers regulate HSPCs using shared or distinct molecular mechanisms. Here, we generate a comprehensive and generalizable cell annotation method (HemaScribe) and a refined quantitative model of hematopoietic differentiation (HemaScape) using single cell RNA sequencing (scRNA-seq) of HSPCs, which we apply to a broad range of EM modalities. We uncover multiple strategies to enhance myelopoiesis acting at different levels of the HSPC hierarchy, which are associated with both unique and shared transcriptional response modules. In particular, we identify a myeloid progenitor-based module of EM engagement across diverse inflammatory challenges, which informs outcome in adult and pediatric human acute myeloid leukemia. Collectively, our work illuminates fundamental regulatory mechanisms in hematopoietic regeneration that have direct translational applications in disease contexts.

**HIGHLIGHTS:**

- New HemaScribe method for hematopoietic progenitor annotation in scRNA-seq datasets
- Different emergency myelopoiesis (EM) inducers act at distinct hematopoiesis levels
- Unique and shared transcriptional response modules enacted by different EM inducers
- A myeloid progenitor EM module informs outcome in acute myeloid leukemia

**eTOC BLURB:** Swann et al. conduct comparative analysis of single cell RNA sequencing data from multiple emergency myelopoiesis models, finding that different perturbations act at various levels of the hematopoietic hierarchy and recruit distinct sets of molecular mechanisms to enhance myelopoiesis. In particular, they identify a conserved myeloid progenitor-based activation module across multiple disease conditions, which informs outcome in human acute myeloid leukemia.

**GRAPHICAL ABSTRACT:** 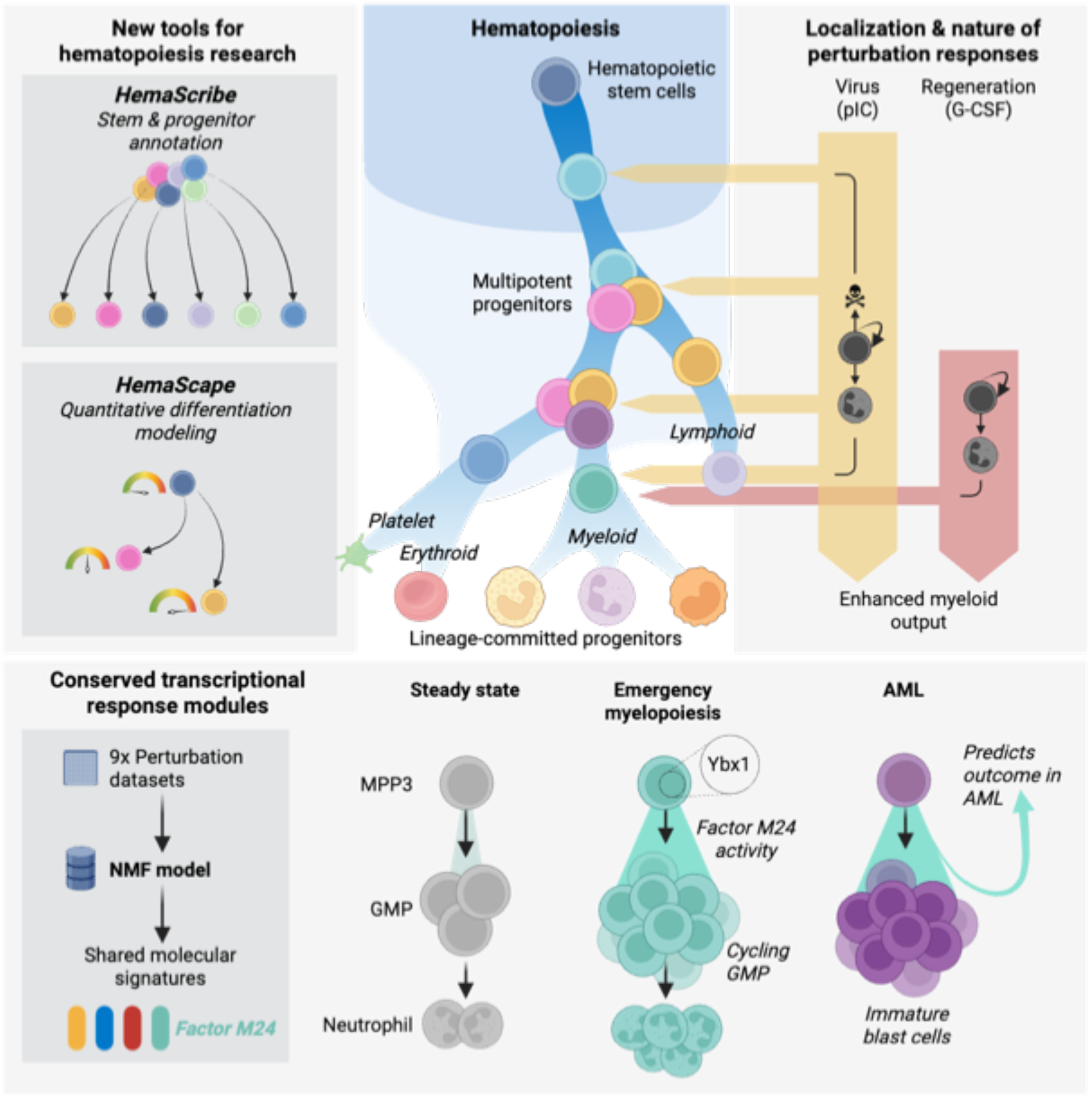

## INTRODUCTION

Hematopoiesis is highly adaptable to demand, as occurs with increased production of mature myeloid cells during infections, chronic inflammation, cancer, and medical interventions like chemotherapy and radiotherapy^1^. In response to these stressors, hematopoietic stem and progenitor cells (HSPC) located in the bone marrow (BM) niche recruit an array of cellular and molecular mechanisms, collectively termed emergency myelopoiesis (EM) pathways, which coordinate rapid production of required myeloid cells^2^. These EM pathways are normally terminated upon resolution of the initiating insult, but there is growing evidence of their persistent or inappropriate activation in aging, cancer, and chronic inflammatory diseases, rendering them important potential targets for therapy^1,2^.

A wide range of mechanisms is important for EM pathway engagement, including increased proliferation^3^, altered cell death^4^, metabolic remodeling^5^, and activation of molecular mechanisms that accelerate myeloid differentiation^6,7^. Among the latter, exposure to interleukin (IL)-1β or tumor necrosis factor (TNF)-α results in precocious activation of the myeloid transcription factor (TF) PU.1 in hematopoietic stem cells (HSC), hence driving enhanced myeloid commitment at the peak of the HSPC hierarchy^8,9^. In the downstream multipotent progenitor (MPP) compartment, the inflammatory cytokine IL-6 can reprogram normally lymphoid-biased MPP4 towards a myeloid fate, which is particularly important during the development of myeloid malignancies^10,11^, while an expanded EM-associated FcγR^+^ subset of myeloid-biased MPP3 undergoes accelerated differentiation toward the granulocyte macrophage progenitor (GMP) stage in conditions of neutrophil depletion or inflammatory stimulation^12^. Moreover, downstream of MPPs, both regenerative and inflammatory stimuli can activate molecular circuits in lineage-committed GMPs that permit transient self-renewal and expansion, leading to the emergence of GMP clusters in the BM niche that act as critical sites of neutrophil amplification in both regenerative and malignant contexts^13^. Collectively, these examples illustrate how HSPC differentiation landscapes can be altered in conditions of enhanced myelopoiesis and highlight the diversity of molecular mechanisms available to the hematopoietic system to respond to increased demand for mature myeloid cells.

Despite this general understanding of EM pathways, it remains unclear whether all EM inducers alter activity at every level of the myelopoietic hierarchy, or if different perturbations exert focused effects on specific populations. This question has become particularly relevant since several murine lineage tracing studies have suggested that HSCs are largely dispensable for EM engagement in response to sepsis or inflammation, inferring that both steady state and regenerative myelopoiesis are solely dependent on MPPs and their progeny^14,15^. It is also currently unknown whether each perturbation activates a unique set of molecular mechanisms in HSPCs, which would be more effective in resolving the initial insult but could carry a high evolutionary cost in encoding and transmitting information for each stimulus across cells and generations. Conversely, we hypothesize that diverse insults will recruit shared effector modules that would represent a more efficient means to encode responses to stimuli, as well as being attractive therapeutic targets for mitigating maladaptive EM responses. While numerous studies, including ours, have dissected the detailed mechanisms observed with single perturbations, there have been few attempts to search for shared modules across biological conditions, or to evaluate their possible clinical significance.

Here, we leveraged the power of our extensive phenotypic and molecular characterization of 9 different perturbations driving EM engagement and covering a range of regenerative, inflammatory, and developmental conditions, to ask how different EM inducers remodel the HSPC hierarchy. To match cell identification and differentiation stages in these distinct single cell RNA sequencing (scRNA-seq) datasets, we first developed new tools for comprehensive cell annotation (HemaScribe) and quantitative trajectory modeling (HemaScape) of HSPC responses. Using these approaches, we demonstrate that while different EM stimuli have distinct effects on cellular targets and preferred downstream molecular mechanisms, they often share transcriptional effector modules with consistent effects in demand-adapted hematopoiesis. In particular, we identify a myeloid progenitor-based EM module that is upregulated in multiple inflammatory and regenerative conditions and has significant prognostic value in human acute myeloid leukemia (AML).

## RESULTS

### Consistent and high-resolution annotation of HSPCs in scRNA-seq datasets

To understand how different perturbations remodel the murine hematopoietic system, we first addressed two major technical challenges in the analysis of scRNA-seq HSPC datasets: (1) how to annotate the same cell types consistently and with high resolution across samples, and (2) how to reconstruct the topology of normal hematopoietic differentiation and quantify its disruption. To overcome the first challenge, we created a new annotation approach called “HemaScribe” (**Fig. 1A**) by first curating bulk RNA sequencing profiles for purified BM cell types from the ImmGen and Haemopedia repositories^16–18^ (**Table S1**) to score cells according to their predicted identity (broad classifier) (**Fig 1B**). This approach was highly effective for mature and differentiated cells in whole BM datasets (**Fig. S1A**) but did not provide sufficient resolution for HSPCs (**Fig. 1B**), particularly for different MPP and lineage-committed progenitor subsets that have undergone extensive functional validation in previous studies^12,19–21^ (**Fig. S1B**). To remedy this, we created a reference dataset by FACS-sorting 10 different HSPC populations (i.e., HSC, short-term [ST]-HSC, MPP2, FcγR positive [FR^+^] and negative [FR^-^] MPP3, MPP4, common lymphoid progenitor [CLP], megakaryocyte progenitor [MkP], erythroid progenitor [EryP], and GMP) before labelling each cell type with different oligo-hash antibodies and pooling them for sequencing (**Fig. S1B,C**). We used this dataset to train a classifier based on anchor integration and label transfer (fine classifier), which achieved high accuracy in reproducing the ground truth cell type represented by oligo hashtags (**Fig. 1C,D; Fig. S1D**). Interestingly, the lowest recall was observed for the most transitional cell types, particularly for MPP2 and the FR^+^ MPP3 subset that we recently described as intermediate between FR^-^ MPP3 and GMP^12^ (**Fig. 1E**). Moreover, our system produced stable results when applied repeatedly to bootstrap samples of the same FACS-sorted Lin^-^/c-Kit^+^ (LK) and Lin^-^/Sca-1^+^/c-Kit^+^ (LSK) scRNA-seq datasets (LK: mean Jaccard similarity of 89.7%, S.D. 8.8; LSK: mean Jaccard similarity of 89.7%, S.D. 8.0), with the lowest similarity indices seen again for the most transitional cell types (FR^+^ MPP3: mean Jaccard similarity of 70.7%, S.D. 10.3) (**Fig. S1B**, **S2A**). We further increased the annotation resolution of the large GMP population using bulk RNA sequencing references for subgroups of granulocyte progenitors (GP), common monocyte progenitors (cMoP), and multilineage GMP (mGMP) that generate either GP or cMoP^22^ (**Fig. S2B,C**). We then applied the complete classifier to any cell labelled as “HSPC” in the bulk classifier to produce our “HemaScribe” annotation (**Fig. 1F; Table S2**), which yielded predicted cell types in LK/LSK scRNA-seq datasets at similar frequencies to those observed by flow cytometry (**Fig. 1G**). To validate our approach, we compared our combined annotations to reference datasets generated by bulk or single cell CITE-seq approaches^12,23,24^, finding high levels of concordance between all these resources (adjusted Rand index 0.84 for bulk samples; 0.70 for CITE-seq) (**Fig. S2D-F**). Lastly, to illustrate how HemaScribe provides greater resolution in HSPC annotation than previous approaches, we re-annotated a published dataset of Lin^-^ BM cells^25^ (**Fig. S3A**), finding that true HSCs were present only in cluster “HSC-1”, whereas “HSC-2” and “HSC-3” contained chiefly MPPs, and that “intermediate myeloid progenitors” (IMP) were primarily cMoP (**Fig. S3B**). To extend the applicability of our classification system to HSPCs and hematopoietic cells residing in extramedullary tissues, we also incorporated a hematopoietic score based on expression of 17 marker genes (**Fig. S3C; Table S3**), which effectively discriminates non-hematopoietic cell clusters in liver and lung datasets^26^, or in BM samples that include stromal cells^27^ (**Fig. S3D,E**). The hematopoietic score and the broad/fine classifiers are now available to the scientific community as the R package “HemaScribe” (**Resource Availability**), with the reference data for the oligo-hashed HSPC populations accessible on an interactive website (**Resource Availability**). Collectively, this work provides a consistent approach to annotate BM datasets, achieving unprecedented resolution in distinguishing HSPC subgroups, and linking cell classification of scRNA-seq data to flow cytometric definitions, which permits coordinated molecular analysis and prospective isolation of the same cell types.

**Figure 1.**
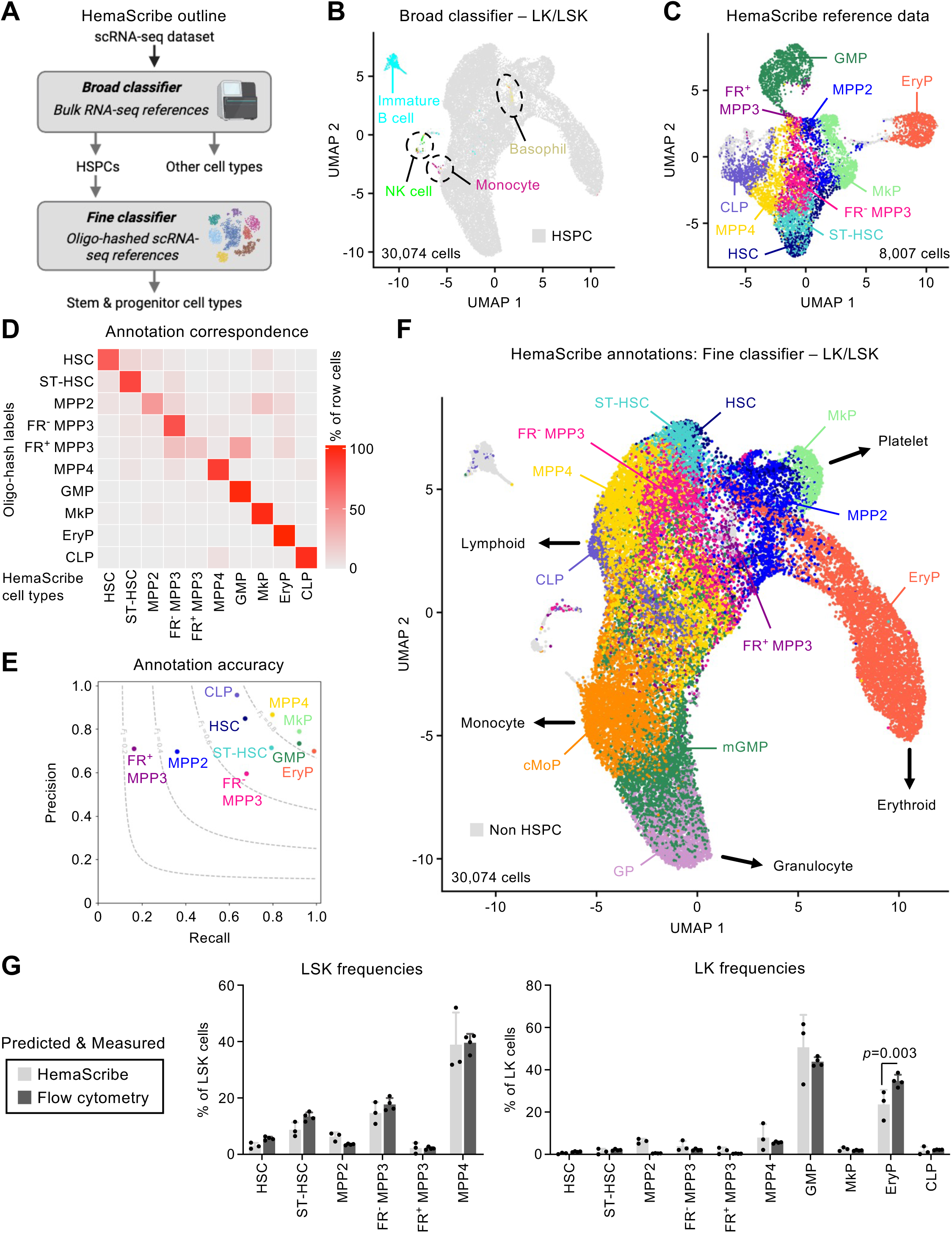
HemaScribe provides consistent annotation of cell types. (**A**) Outline for classification of cell types using bulk and single cell RNA sequencing (scRNA-seq) references. HSPC: hematopoietic stem and progenitor cell. (**B**) Uniform manifold and approximation projection (UMAP) showing annotation of 10X scRNA-seq dataset from Lin^-^/c-Kit^+^ (LK) and Lin^-^/Sca-1^+^/c-Kit^+^ (LSK) BM cells. (**C**) UMAP showing 10 different sorted HSPC populations annotated according to HemaScribe fine classifier results. CLP: common lymphoid progenitor; EryP: erythroid progenitor; FR^-/+^ MPP3: FcγR^-/+^ multipotent progenitor 3; GMP: granulocyte macrophage progenitor; HSC: hematopoietic stem cell; MkP: megakaryocyte progenitor; MPP: multipotent progenitor; ST-HSC: short-term HSC. (**D**) Heatmap showing correspondence between hashtag labels and HemaScribe fine classifier results for indicated cell types. (**E**) Precision and recall for fine classifier in indicated cell types. (**F**) UMAP showing annotation of LK/LSK scRNA-seq data using HemaScribe fine classifier and GMP subclassification. cMoP: common monocyte progenitor; GP: granulocyte progenitor; mGMP: multilineage GMP. (**G**) Bar charts showing frequency of indicated cell types among LSK (left) or LK (right) cells, derived from either flow cytometry or by annotation of scRNA-seq datasets using the HemaScribe fine classifier. Points represent individual mice (flow cytometry) or biological replicates of scRNA-seq with mean ± S.D.; *P. values*, two-way ANOVA with Sidak’s *post hoc* test. See also Figures S1, S2, and S3.

### A unified topology for hematopoietic differentiation

To overcome the second challenge of understanding how HSPCs differentiate at steady state, we developed the “HemaScape” tree model to visualize the topology of hematopoiesis (**Fig. 2A**). Using our reference oligo-hashed scRNA-seq dataset, we employed a dimensionality reduction method (elastic embedding [EE])^28^ to capture both local and global continuous structure, before locating regions of high cell density taken to represent stable cell states using the DensityPath algorithm^29^. To delineate connections among these density clusters, we calculated minimum spanning trees that can represent underlying differentiation trajectories (**Fig. 2B**), and we used the same density clusters as nodes for partition-based graph abstraction (PAGA) analysis^30^ to quantify connectivity estimates among nodes (**Fig. 2C; Table S4**). We combined the branching structure and connectivity values into a HemaScape differentiation tree that depicts the arrangement of major HSPC branches (**Fig. 2D**) and which is directly cross-referenced with our HemaScribe cell type annotation (**Fig. S4A**). As validation for this model, we compared the HemaScape tree with an alternative structure inferred from rates of propagation of a fluorescent label originating in HSCs across a longitudinal time course in *Hoxb5-Cre:R26LSL-tdT* mice^31^ (**Fig. 2E**). To evaluate the similarity of the dendrogram between these models, we first randomly permuted the edges of the HemaScape tree 10,000 times and found that our solution significantly resembled the structure outlined by label propagation data (Pearson r=0.41, *P*=0.04) (**Fig. 2F**), indicating that both models support the same general arrangement of lineage branches in hematopoiesis. Next, we assessed the extent to which each model captured transcriptional cell states by computing the distribution of cell type labels for each node across multiple independent replicates, finding that nodes in the HemaScape tree showed greater correspondence with cell types than those in the label propagation dataset (**Fig. 2G**). By comparing limbs of the HemaScape tree, we also captured differentially regulated SCENIC TF regulons^32^ at major branch points, which included many known determinants of commitment to erythropoiesis (Gata1)^33^, myelopoiesis (C/EBPα)^34^, and lymphopoiesis (Ets1, Tcf4)^35^ (**Fig. S4B**). Finally, to understand the cellular behavior of different nodes in the HemaScape tree, we used a simplified version of our proposed structure that condensed smaller nodes in lineage-committed branches (**Fig. 2H**). We estimated cell death rates by direct measurement of ApoTracker green staining (**Fig. S4C,D**) and proliferation rates by calibrating a gene expression score of cell cycle activity against HSPCs identified in the Fucci2 cell cycle reporter mouse line^36^ (**Fig. S5A,B; Table S5**). Using these parameters combined with node connectivity values from PAGA and estimates for node sizes from their natural frequencies in steady state LK datasets, we then implemented an iterative random walk approach to estimate the “differentiation rate” for each node, representing the likelihood that cells located in any node would progress to another. Doing so, we inferred higher proliferation and differentiation rates in downstream lineage-committed progenitor nodes compared to HSC/MPPs (**Fig. 2H; Table S6**), and especially in the megakaryocyte/erythroid (MegE) branches, which aligns with patterns observed from label propagation data^31^ (**Fig. 2E**). Moreover, using Palantir^37^ (**Fig. S5C,D; Table S7**), we confirmed a tendency for greater transcriptional entropy of immature HSC/MPPs compared to lineage-committed progenitors in the erythroid or myeloid lineages (**Fig. 2H**), which approximates the greater multipotency of HSPCs at the peak of the hematopoietic hierarchy. This approach also allowed us to confirm the terminal state probabilities of different MPP subsets inferred previously by transplantation^21^ and barcode sharing data^38^, with MPP2 showing the greatest bias towards MegE cells, FR^+^ MPP3 towards myeloid fate, and MPP4 towards lymphoid fate (**Fig. S5E**). Taken together, our findings outline a quantitative model for hematopoietic differentiation at steady state that recapitulates numerous experimental observations, produces better capture of transcriptional states, and enables comparison to perturbed conditions. We provide functions to map query datasets to our EE embedding and assign HemaScape nodes in the R package “HemaScribe” (**Resource Availability**).

**Figure 2.**
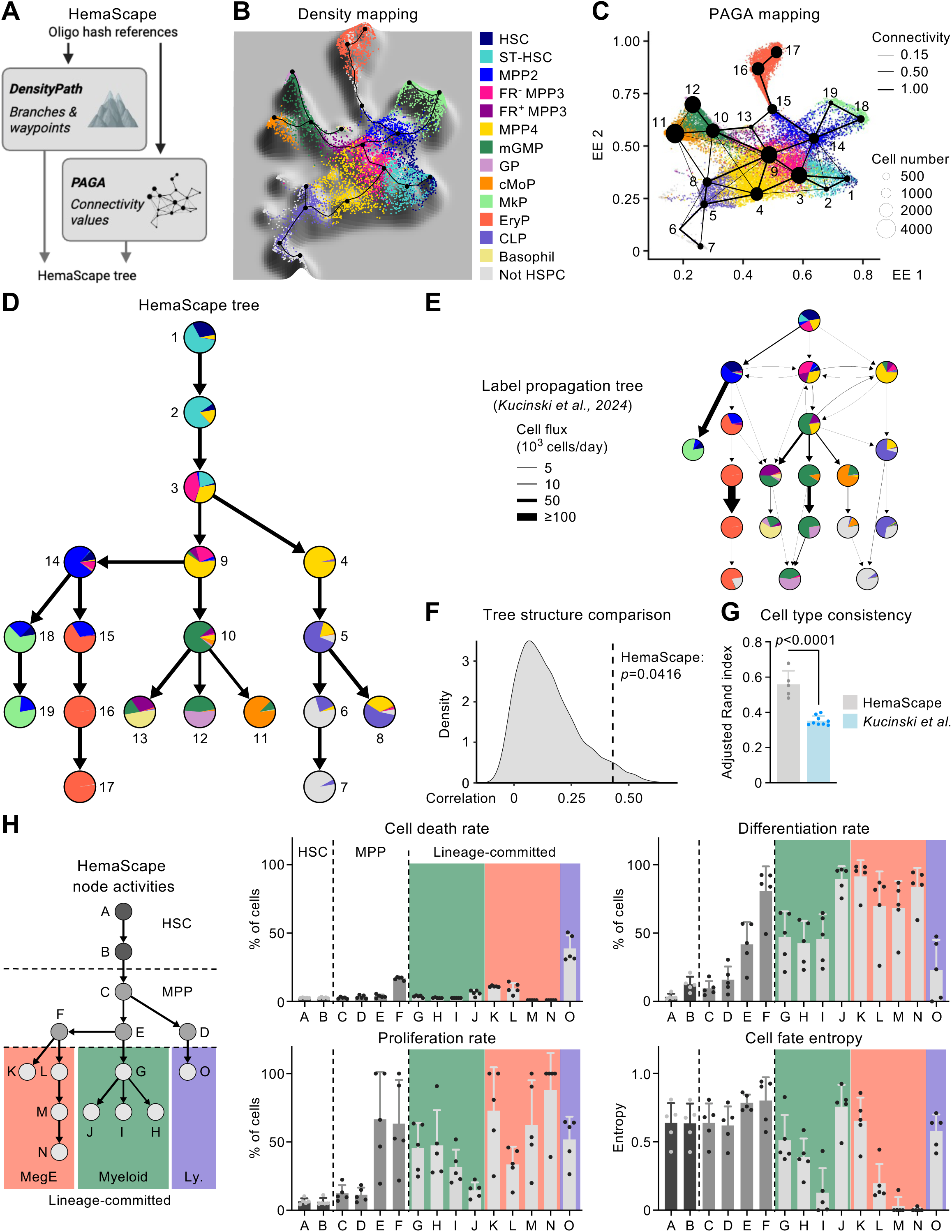
HemaScape topological landscape for hematopoietic differentiation. (**A**) Outline showing how DensityPath was used to identify density clusters, which were connected by minimum spanning trees. Density clusters were also used as input for partition-based graph abstraction (PAGA) analysis, and these measures were combined to infer the HemaScape tree. (**B**) Density landscape showing density clusters (black points) and minimum spanning tree (black lines), annotated by cell type. (**C**) Elastic embedding (EE) showing PAGA nodes derived from density clusters. (**D**) HemaScape tree structure inferred from density-based branching structure, with edge thickness determined by PAGA connectivity, and nodes colored by HemaScribe composition. (**E**) Tree structure proposed in previous publication of label propagation data, where edge thickness indicates differentiation flux and nodes colored by HemaScribe annotation. (**F**) Histogram showing results of perturbation of HemaScape tree and Pearson correlation with tree structure in (E). (**G**) Bar graph showing adjusted Rand index comparing HemaScribe cell type composition between nodes in HemaScape tree and tree in (E). Points represent independent scRNA-seq replicates with mean ± S.D.; *P.* value from Student’s t-test. (**H**) Simplified HemaScape tree structure (left) used for estimation of node size, proliferation, cell death, connectivity values, and differentiation rates. Bar charts (right) show calculated cell death rates, inferred proliferation rates, inferred differentiation rates, and Palantir entropy for nodes shown in tree structure. MegE: megakaryocyte/erythroid; Ly: lymphoid. Points represent biological replicates of LK scRNA-seq datasets (n=5), with mean ± S.D.; *P. value* from permutation testing. See also Figures S4 and S5.

### Different perturbations affect distinct levels of the hematopoietic system

We next compared scRNA-seq datasets from several EM perturbations to evaluate changes in HSPC landscape, incorporating 7 models representing inflammation, regeneration, and responses to pathogen-derived products. Inflammation was induced by 7 (short-term) or 20 (chronic) daily injections of IL-1β^8,39^, regeneration by 2 or 4 daily injections of granulocyte colony stimulating factor (G-CSF) or occurring 8 days after administration of the cytotoxic drug 5-fluorouracil (5FU)^13^, and response to pathogen-derived products was studied at 16 hours after lipopolysaccharide (LPS) injection^23^ or 3 days after polyinosinic:polycytidylic acid (pIC) injection^40^ (**Fig. S6A**). As controls, we also included a published dataset from mice exposed to the hormone erythropoietin (EPO) for 2 days^41^ to drive increased output in the erythroid lineage, and published datasets from embryonic day 18.5 (E18.5) fetal^23^ and 24-month-old aged^42^ mice, in which differences in HSPC frequency have been described previously. We confirmed that many EM perturbations caused increases in peripheral blood neutrophil counts, particularly with IL-1 and G-CSF treatments, whereas 5FU caused the expected depletion of mature neutrophils at this timepoint^13^ (**Fig. S6B**). To evaluate quantitative changes in cell populations, we mapped LK scRNA-seq datasets to our reference embedding (**Fig. S6C**), applied our annotation approach to each dataset (**Fig. 3A; Table S3**), and compared cell frequencies using single cell differential composition analysis (scDC)^43^. Importantly, by comparing our classifier trained on steady state populations with a similarly generated complementary algorithm trained on 5FU-treated HSPC populations, we validated that our standard cell type identification approach performed well in perturbation conditions, despite major changes in gene and surface marker expression (**Fig. S7A-C**). Comparing our HemaScribe annotation results, we observed marked differences in the effect of different EM perturbations on cell type distributions in the hematopoietic system. Consistent with previous findings^13^, 5FU caused massive expansion of HSCs (14.6x) and MPPs at the expense of lineage-committed progenitors in all lineages (**Fig. S7D**). In contrast, the pathogen-derived products pIC and LPS caused more subtle changes in MPPs, with pIC also increasing the frequency of mGMP (2.2x) (**Fig. 3B; Fig. S7E**). Administration of G-CSF or IL-1 had little impact on HSC and MPP frequency but caused dramatic expansion of GMPs, especially mGMP and GP (2.4x and 2.3x at day 4 of G-CSF treatment, respectively) at the expense of EryP (**Fig. 3C; Fig. S7F**), whereas EPO had the opposite effect of increasing EryP (1.6x) and decreasing GMP subsets (**Fig. S7G**). HemaScribe annotation of fetal and aged datasets also confirmed previous findings^23,42^, with expanded EryP (2.0x) at the expense of GMPs in E18.5 embryos, and a dramatically enlarged HSC compartment (19.2x) with increased mGMP (1.9x) and decreased EryP in old mice compared to their respective 3-month-old adult controls (**Fig. S7H,I**). Taken together, these results indicate that different EM perturbations and developmental states produce distinct patterns of change in HSPC frequency that target different levels of the hematopoietic hierarchy.

**Figure 3.**
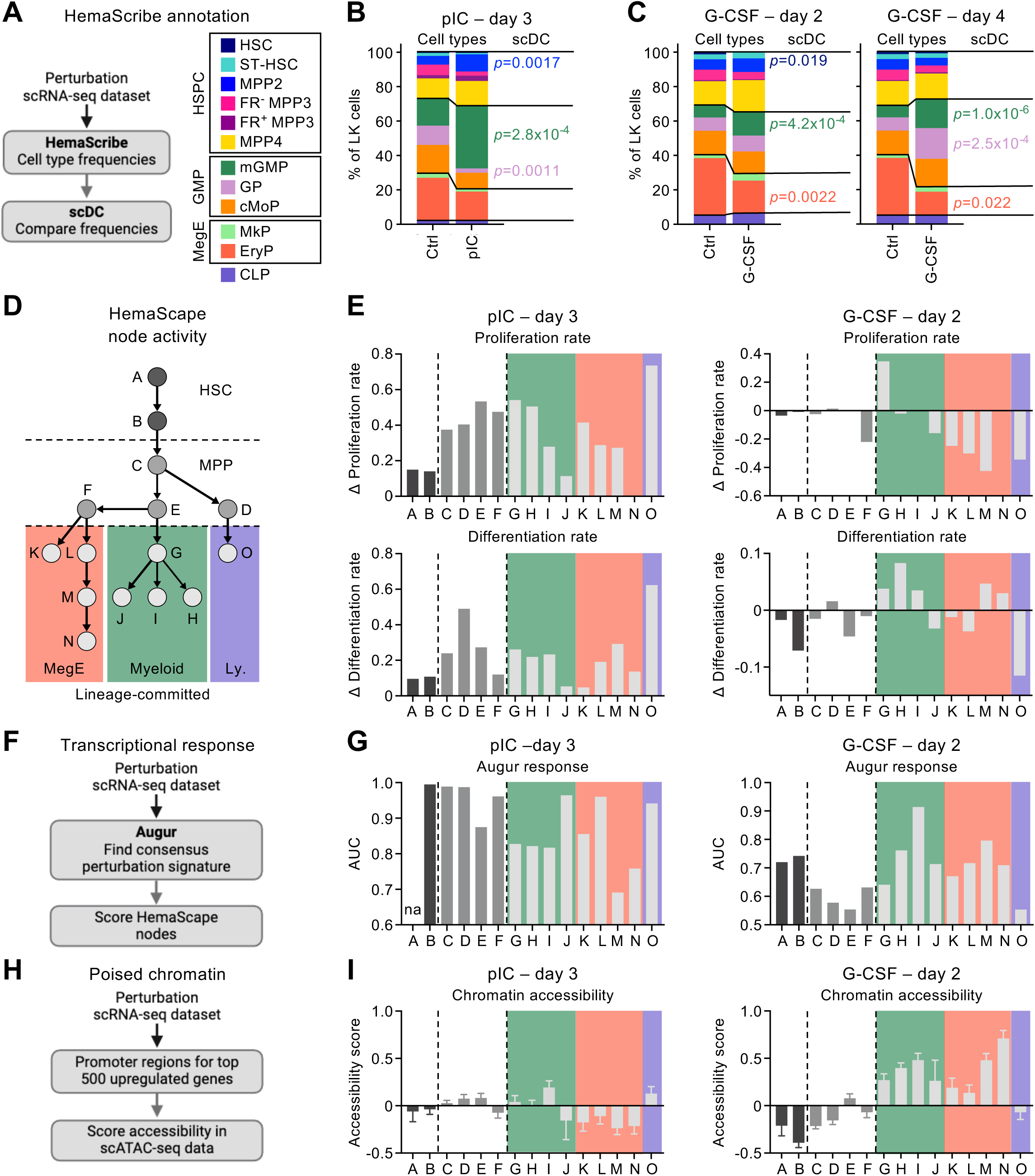
Emergency myelopoiesis perturbations act at different levels of the hematopoietic hierarchy. (**A**) Outline and key for HemaScribe annotation of perturbation datasets and matched controls. (**B-C**) Stacked bar charts showing frequency of indicated cell types in LK datasets for (B) 3-day polyinosinic:polycytidylic acid (pIC) exposure and (C) 2- or 4-day granulocyte colony stimulating factor (G-CSF) treatment; *P. values* derived from general linear models using single cell differential composition (scDC). (**D**) Simplified HemaScape tree used as template for estimating node activities. (**E**) Bar charts showing inferred changes in proliferation (top) and differentiation rate (bottom) for HemaScape nodes from transcriptional inference and random walk models compared to matched controls for 3-day pIC and 2-day G-CSF datasets. (**F**) Outline for assessment of transcriptional responses in HemaScape nodes using Augur. (**G**) Bar charts showing predicted transcriptional response expressed as area under curve (AUC) from Augur. *na* indicates insufficient cells for calculation. (**H**) Outline for assessment of epigenetic poising for response to perturbations. (**I**) Bar charts showing mean difference in accessibility score for promoters of genes upregulated by indicated perturbations compared to promoters of random set of genes. Bars show mean difference ± S.E.M. See also Figures S6, S7, and S8.

To understand the cellular processes driving the observed quantitative changes, we recreated estimates of HemaScape node activity for each perturbation, just as we had implemented for steady-state hematopoiesis (**Fig. 3D; Table S8,S9**), focusing on the effects of pIC and G-CSF as different classes of EM driver. Our model predicted that pIC would broadly increase proliferation and differentiation rates in MPP (C-F) and downstream myeloid progenitor nodes (G-J), whereas G-CSF would instead cause a focused increase in proliferation only in the mGMP-enriched node G and differentiation in most of myeloid progenitor nodes (G-I) but not MPP nodes (**Fig. 3E**). Interestingly, these predictions based only on quantitative data were often mirrored by evaluation of the same nodes using Augur^44^, which measures the strength of the transcriptional response to perturbation (**Fig. 3F,G**). This revealed little bias for pIC, with the greatest transcriptional response in ST-HSC and MPP nodes (B-D), whereas G-CSF caused the greatest effect in myeloid progenitor nodes (H-I) (**Fig. 3G**). Interestingly, LPS produced a pattern similar to pIC, which was dominated by increased proliferation, differentiation, and transcriptional response in MPPs, whereas 5FU caused a more global response that extended also to lineage-committed progenitors (**Fig. S8A**), consistent with the broad regenerative response to this drug. To understand if the predicted differences between pIC and G-CSF were hardwired in the epigenetic state of different cell types, we exploited a combined LK/LSK multiome dataset^23^ containing both RNA and chromatin accessibility data, which we annotated using HemaScribe on the RNA assay (**Fig. S8B**). We then generated signatures based on the genes upregulated by each perturbation and scored each cell type for accessibility of those regions (**Fig. 3H**). Strikingly, committed myeloid progenitor nodes (G-J) showed increased resting accessibility for genes upregulated by G-CSF compared to a random sample of genes, whereas no cell type was particularly enriched for the gene signature induced by pIC (**Fig. 3I**). Collectively, these data suggest that myeloid progenitors are epigenetically poised and competent to respond to G-CSF, which induces localized EM transcriptional, proliferation, and differentiation responses to increase myeloid cell production, without affecting upstream HSCs and MPPs. In contrast, no specific cell type appears to be epigenetically predisposed to respond to pIC, which probably accounts for the broader range of EM transcriptional and differentiation responses in this context, including activation of early HSPCs.

### Emergency myelopoiesis alters differentiation pathways

Next, we evaluated whether our HemaScape model could predict the engagement of new differentiation links, such as MPP reprograming or bypass mechanisms that have been described recently in experimental studies^2^. To do so, we calculated PAGA connectivities for each perturbation and its matched control sample and filtered for links showing changes of 0.15 or more (**Fig. 4A; Table S8**). We first evaluated patterns of connectivity involving early HSPCs, finding that, among all perturbations, only pIC and LPS increased links between HSC and either MPP2 or mixed MPP nodes (**Fig. 4B; Fig. S9A**), supporting the results of our node activity estimates. Moreover, we found that both pIC and LPS also specifically increased connectivity between HSC and megakaryocyte progenitor (MkP) nodes (**Fig. 4C; Fig. S9B**), consistent with the reported HSC-MkP bypass mechanism^45,46^, for which pIC and downstream type I interferon (IFN) responses are major drivers^47,48^. To test this association directly, we isolated HSCs from *β-actin-Gfp* mice^49^ and transplanted them into lethally irradiated recipients (**Fig. 4D**). Starting from the day after transplant, we injected the recipients with either pIC or G-CSF according to the same dose and schedule as for our molecular analyses (**Fig. S6A**), finding that only pIC boosted the production of GFP^+^ platelets in peripheral blood over the following two weeks (**Fig. 4D; Fig. S9C**). At the molecular level, SCENIC analysis of TF regulons in pIC-exposed HSCs showed upregulation of factors with essential roles in specifying the megakaryocyte lineage, like Fli1 and Runx3^[50,51]^ (**Fig. 4E**), suggesting pIC stabilizes the MkP bypass pathway by causing precocious activation of platelet lineage-defining TFs. Finally, we evaluated the connectivity of MPP4-enriched nodes because this population biased towards lymphopoiesis at steady state can be reprogrammed by IL-6^[10]^ or TF balance^6^ to produce myeloid cells upon demand. Among all perturbations, we found that LPS, pIC, and IL-1 enhanced connectivity from MPP4/CLP to mGMP/cMoP nodes, whereas G-CSF and 5FU had no effect (**Fig. 4F; Fig. S9D**). To evaluate the same responses *in vivo*, we transplanted purified MPP4 into sub-lethally irradiated congenic recipients and again started either pIC or G-CSF treatments the day after transplant, for a total of 2 cycles (**Fig. 4G**). In line with our prediction, we found that pIC, but not G-CSF, prolonged the myeloid output of MPP4, especially during the first 2 weeks following transplantation (**Fig. 4G; Fig. S9E**). SCENIC analysis of TF regulons in pIC-exposed MPP4 again revealed upregulation of factors that can drive commitment of MPPs to myeloid fate like C/EBPα and C/EBPβ^6^ (**Fig. 4H**). Accordingly, *in silico* knockouts of C/EBPα and C/EBPβ predicted much greater embedding shifts towards MPPs in pIC-treated GMP than in G-CSF-treated cells (**Fig. S9F**), suggesting a role for pIC-induced differential regulation of C/EBP factors in altering MPP4 behavior. Taken together, these results show that pathogen-derived products, especially pIC, can remodel the entire HSPC hierarchy through MPP reprograming and engagement of differentiation bypass mechanisms, whereas G-CSF acts only by enhancing myeloid progenitor-specific pathways that already operate at steady state (**Fig. S10A**). Moreover, they illustrate the power of our quantitative HemaScape model in predicting the differentiation paths used specifically by each EM inducer, creating a unique EM fingerprint delineated by activation of distinct molecular mechanisms at different levels of the HSPC hierarchy.

**Figure 4.**
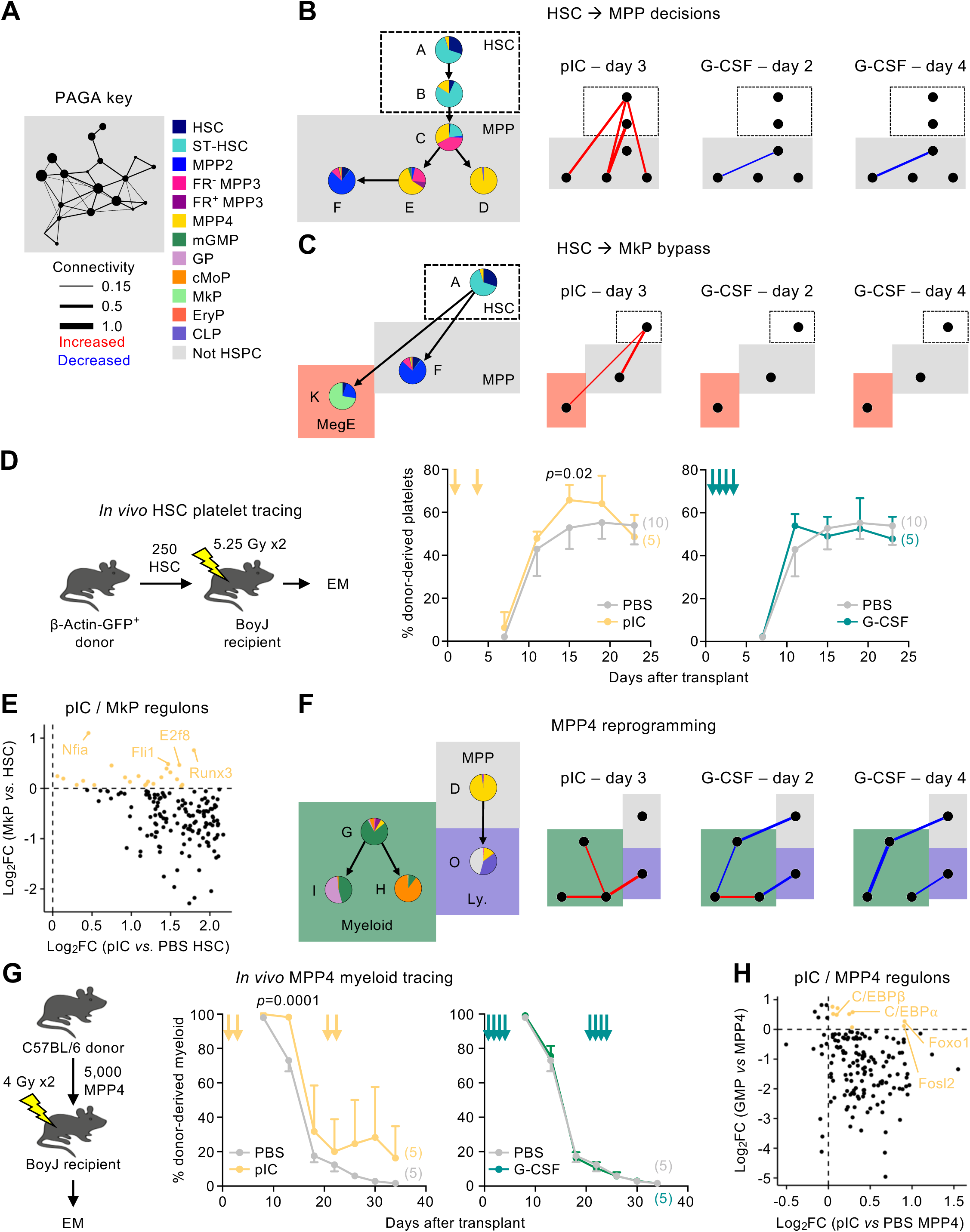
Pathogen-derived stimuli exploit alternative differentiation pathways. (**A**) Outline showing how changes in PAGA connectivity were evaluated in indicated perturbations compared to matched control datasets. (**B**) Outline of HemaScape nodes evaluated for changes in connectivity (left, see Figure 2H), and subplots showing calculated changes for indicated perturbations. (**C**) Outline and PAGA subplots showing changes in connectivity between indicated HemaScape nodes. (**D**) Outline (left) for transplantation of fluorescent HSC into lethally irradiated recipients followed by injection of pIC or G-CSF to stimulate emergency myelopoiesis (EM). Arrows indicate timing of injections. Quantification (right) of platelet chimerism in peripheral blood for recipient mice. Points represent n=5-10 mice per group, with mean ± S.D. and *P. values* from multiple t-tests. (**E**) Scatter plot showing log2 fold change (FC) in SCENIC transcription factor regulons comparing steady state MkP to HSC, against pIC-exposed HSC comparing to control HSC. (**F**) Outline and PAGA subplots showing changes in connectivity among indicated HemaScape nodes. (**G**) Outline (left) showing how MPP4 were transplanted into sublethally irradiated congenic recipients followed by injection of pIC or G-CSF to induce EM. Arrows indicate timing of injections. Quantification (right) of proportion of output of transplanted MPP4 that was myeloid (expressing CD11b) in peripheral blood of recipients. Points represent n=5 mice per group, with mean ± S.D. and *P. values* from multiple t-tests. (**H**) Scatter plot showing log2FC in SCENIC transcription factor regulons comparing steady state GMP to MPP4, against pIC-exposed MPP4 comparing to control MPP4. See also Figures S9 and S10.

### Shared and distinct molecular patterns of hematopoietic activation

Although each perturbation acts at different levels of the HSPC hierarchy, we wondered whether common transcriptional modules could be activated by different EM stimuli to enhance myeloid output. To achieve this, we performed non-negative matrix factorization (NMF) on an integrated dataset containing 9 conditions (5FU day 8, G-CSF day 2 and day 4, IL-1 day 7 and day 20, LPS 16 hours, pIC day 3, aged 24 months, and fetal E18.5) and consisting of 25 scRNA-seq samples (**Fig. 5A; Fig. S10B; Table S10**). Cross-validation suggested a total of 40 factors was appropriate for our dataset (**Fig. S10C**), which we categorized manually into those representing sources of technical or batch-related variation, cell type specific effects, or processes that varied by perturbation (**Fig. S10D; Table S11**). We also established a myeloid pseudotime gradient, beginning with HSC and progressing through density clusters to reach GP and cMoP (**Fig. S10E**). When we evaluated constitutive molecular processes along this gradient, we found a clear succession starting with an HSC-associated murine factor (M7) driven by stemness genes like Hoxa10, progressing through MPP-related factors (M9, M13) associated with Hoxa7 and Sox4 genes, and culminating with myeloid-related factors (M11, M23, M15, M30) associated with well-known myeloid TFs like C/EBPα, IRF8, C/EBPε, and Gfi1 (**Fig. 5B**). Among the factors that varied by perturbation, we selected 4 (M12, M18, M24, and M40) for further consideration. M18 was most active among HSC and ST-HSC, and its top loading genes included those encoding the TFs Runx1, Etv6, and Dach1 (**Fig. 5C**). M18 was specifically associated with acute IL-1 exposure and both pathogen stimuli, LPS and pIC (**Fig. 5C**), and displayed the highest level of conservation across vertebrate species (**Fig. S10F**). Gene ontology (GO) analyses of its loading genes showed enrichment for pathways involved in immune system processes (**Fig. S10G**), and SCENIC regulon scores indicated the greatest association with Tcf12 and Runx1 (**Fig. 5D**). This suggests that M18 is associated with acute activation of early HSPCs, likely driven by Runx1 and associated TFs, consistent with the role of Runx1 in driving differentiation into multiple lineages^52^ and specifically activating PU.1 to promote myelopoiesis^53^. While M12 was also most enriched in HSC and ST-HSC, it was activated in different conditions, particularly by chronic IL-1 and 5FU (**Fig. 5E**). M12 was strongly associated with expression of both Jun and Fos family members in its top loading genes, showed strong correlations with the SCENIC regulons for AP-1 TFs, and was enriched for GO pathways related to cell stimulation (**Fig. 5D; Fig. S10G**). This suggests M12 is activated in chronic inflammatory settings and may promote myeloid differentiation in HSPCs, consistent with the role of Jun as an essential co-factor for PU.1 in myelopoiesis^54^. M40 was also most enriched in HSC and ST-HSC and was strongly associated with LPS and pIC (**Fig. 5F**). Accordingly, its top loading genes contained many IFN-induced genes, including *Ly6a* (Sca-1) and *Irf7*, which were enriched for GO pathways related to viral infection, correlated with key IFN-related TFs like Stat1, Stat2, and Irf7 in SCENIC regulons, and enriched for genes induced by IFNs in immune response enrichment analysis (IREA)^55^ (**Fig. 5D,F; Fig. S10G**). Therefore, M40 explains the type I IFN response induced by toll-like receptor (TLR) ligands like LPS and pIC, which probably accounts for its more widespread activity in the hematopoietic system than either M12 or M18 (**Fig. 5G**). In contrast, M24 was most active in myeloid progenitors like GMP, with activity associated with multiple perturbations including 5FU, G-CSF, pIC, and LPS (**Fig. 5H**). Its top loading genes included metabolic genes (*Ldha*, *Cox7b*) as well as interferon-induced transmembrane proteins (*Ifitm1*, *Ifitm3*) normally more highly expressed in HSCs than GMPs^20^, and these were enriched for GO pathways related to defense and innate immune responses (**Fig. 5H; Fig. S10G**). This suggests that M24 is broadly associated with myeloid activation across perturbations.

**Figure 5.**
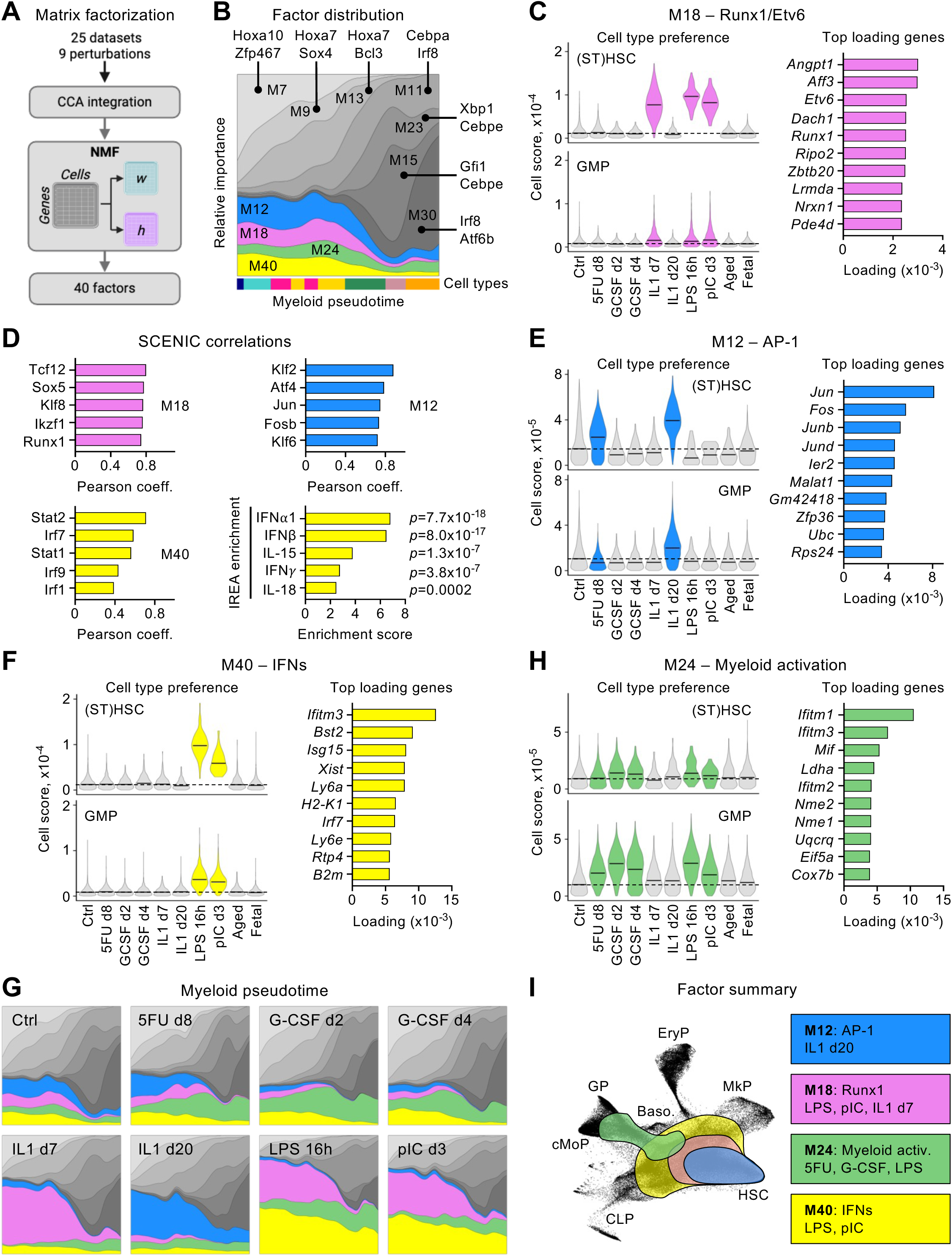
Shared and distinct molecular signatures across emergency myelopoiesis perturbations. (**A**) Outline showing how different EM datasets with integrated by canonical correlation analysis (CCA) before non-negative matrix factorization (NMF), finding k=40 factors. (**B**) Succession of NMF factor activities across myeloid pseudotime, with top SCENIC regulons correlated with each factor. Most abundant cell types in pseudotime bins are shown in colored bar. (**C**) Violin plots (left) showing activity of NMF factor M18 in HSC and ST-HSC (top) or GMP (bottom) across perturbations. Bar charts showing loading scores for top 10 genes for each NMF factor (right). (**D**) Bar charts showing top 5 Pearson correlation coefficients (coeff.) for NMF factor scores and SCENIC regulons for indicated factors, and results of immune response enrichment analysis (IREA) for M40 with *P. values* from two-sided Wilcoxon rank sum test. (**E-F**) Violin and bar charts for indicated NMF factors as outlined in (C). (**G**) Change in relative importance of indicated factors across myeloid pseudotime in indicated perturbations. (**H**) Violin and bar charts for indicated NMF factors as outlined in (C). (**I**) Summary showing major cell types affected by different NMF factors and their key biological attributes. Myeloid activ.: myeloid activation. See also Figures S10 and S11.

Since these factors were identified in datasets representing single biological replicates, we wished to determine whether they could also be observed in independently published resources that used similar *in vivo* EM perturbations, including acute IFN-α treatment over a time course of 72 hours^56^, at 2 hours after pIC and G-CSF injections^57^, and with 5 weeks of chronic IL-1 injection^58^. To achieve this, we used our NMF model to derive scores for each factor in cells of query datasets using non-negative least squares regression before comparing among perturbation conditions (**Fig. S11A; Table S12**). We confirmed that M40 was strongly associated with both type I IFN exposure driven by IFN-α or pIC, representing the most significantly upregulated factor in both settings (**Fig. S11B,C**). Conversely, M12 was the second most upregulated factor with chronic IL-1 (**Fig. S11D**) and, to a lesser extent, with an independently generated time course of 5FU-mediated regeneration, aligning well with our own findings (**Fig. S11E**). We also confirmed strong induction of M18 by pIC but not short-term IFN-α exposure (**Fig. S11B,C**), suggesting this factor could be activated by some other element of TLR signaling. Finally, M24 was more strongly induced in myeloid-committed progenitors than HSCs in an independent G-CSF treatment dataset, as we observed in our own data (**Fig. S11C**). Collectively, these results show that several distinct molecular effector modules can be activated in HSPCs by different groups of perturbations (**Fig. 5I**) in a manner that is highly repeatable across independent studies and in ways that are likely to have important functional consequences for downstream EM responses.

### A myeloid activation factor is conserved in inflammatory and neoplastic diseases

The NMF factor M24 was increased in myeloid-committed progenitors across multiple conditions, making it an attractive core EM module for further study. In the context of 5FU exposure, we confirmed that M24 was among the most differentially increased molecular processes at day 8, representing the time of maximal HSPC activation, in both myeloid-biased MPP3 and GMP (**Fig. 6A,B; Fig. S12A**). Moreover, analysis of published bulk RNA-seq of BM GMP (**Table S13**) from studies of murine myocardial infarction^59^, inflammatory colitis induced with dextran sodium sulfate (DSS)^60^, inflammatory spondyloarthritis^61^, and sepsis caused by LPS exposure^62^ revealed significant enrichment for the top 100 loading genes of M24 in all settings (**Fig. 6C**), highlighting its commonality across inflammatory disease models. M24 top loading genes were significantly enriched for those induced by myeloid cytokines like macrophage colony stimulating factor (M-CSF) and granulocyte macrophage colony stimulating factor (GM-CSF) in IREA analyses (**Fig. 6D**), as well as for genes associated with high output HSCs^63^ (**Fig. S12B**), consistent with M24 being a driver of myeloid regeneration. Exploring possible upstream regulators, we found that M24 correlated highly with the SCENIC regulons for Erf, Ybx1, and Nfia (**Fig. 6E**). To understand these results in the wider context of molecular regulation of myelopoiesis, we performed *in silico* knockout screens of 33 TFs with known roles in myelopoiesis (**Table S14**) using CellOracle^64^ in scRNA-seq datasets, comparing day 8 5FU with day 20 IL-1, with the latter representing a situation lacking increased M24 activity (**Fig. 6F**). Constitutive myeloid factors like C/EBPα were equally important in both conditions, whereas PU.1, Myc, and IRF8 were more important for chronic IL-1 exposure, consistent with the known role of the IL-1-induced NF-κB pathway in activating PU.1^[8]^. Conversely, other factors were more important for 5FU treatment, including C/EBPβ, Xbp1, and Ybx1. Based on these results, we identified the Y-box TF and cold shock protein Ybx1 as a putative regulator of M24. However, the role that this multifunctional protein^65^, which regulates a range of processes important for cell differentiation as well as binding RNA to influence mRNA stability, splicing, and translation, could play in myeloid regeneration is currently unknown. To investigate this, we first assessed Ybx1 protein levels by flow cytometry in myeloid progenitors, confirming significant increases in day 8 5FU-treated MPP3, though not in GMP, with no change in day 7 IL-1-treated MPP3 (**Fig. 6G**). We then performed *ex vivo* liquid culture of GMP isolated from either day 8 5FU-treated, day 7 IL-1-treated, or matching PBS-treated control mice, and expanded them in the presence or absence of the Ybx1 inhibitor (Ybx1i) SU056^[66]^ or PU.1 inhibitor (PU.1i) DB2313^[67]^ (**Fig. 6H**). Consistent with M24 activation, 5FU-treated GMP expanded significantly faster than controls or IL-1-treated GMP. In this context, Ybx1i exposure completely abrogated the expansion of 5FU-exposed GMP, reducing it to the level of control GMP. Importantly, Ybx1i had no significant effect on control GMP or IL-1-treated GMP, nor did PU.1i exposure suppress the expansion of 5FU-treated GMP, demonstrating the specificity of this effect (**Fig. 6H**). Collectively, these data emphasize the broad activity of M24 in settings of myeloid regeneration and identify Ybx1 as a putative and targetable functional driver of this molecular process.

**Figure 6.**
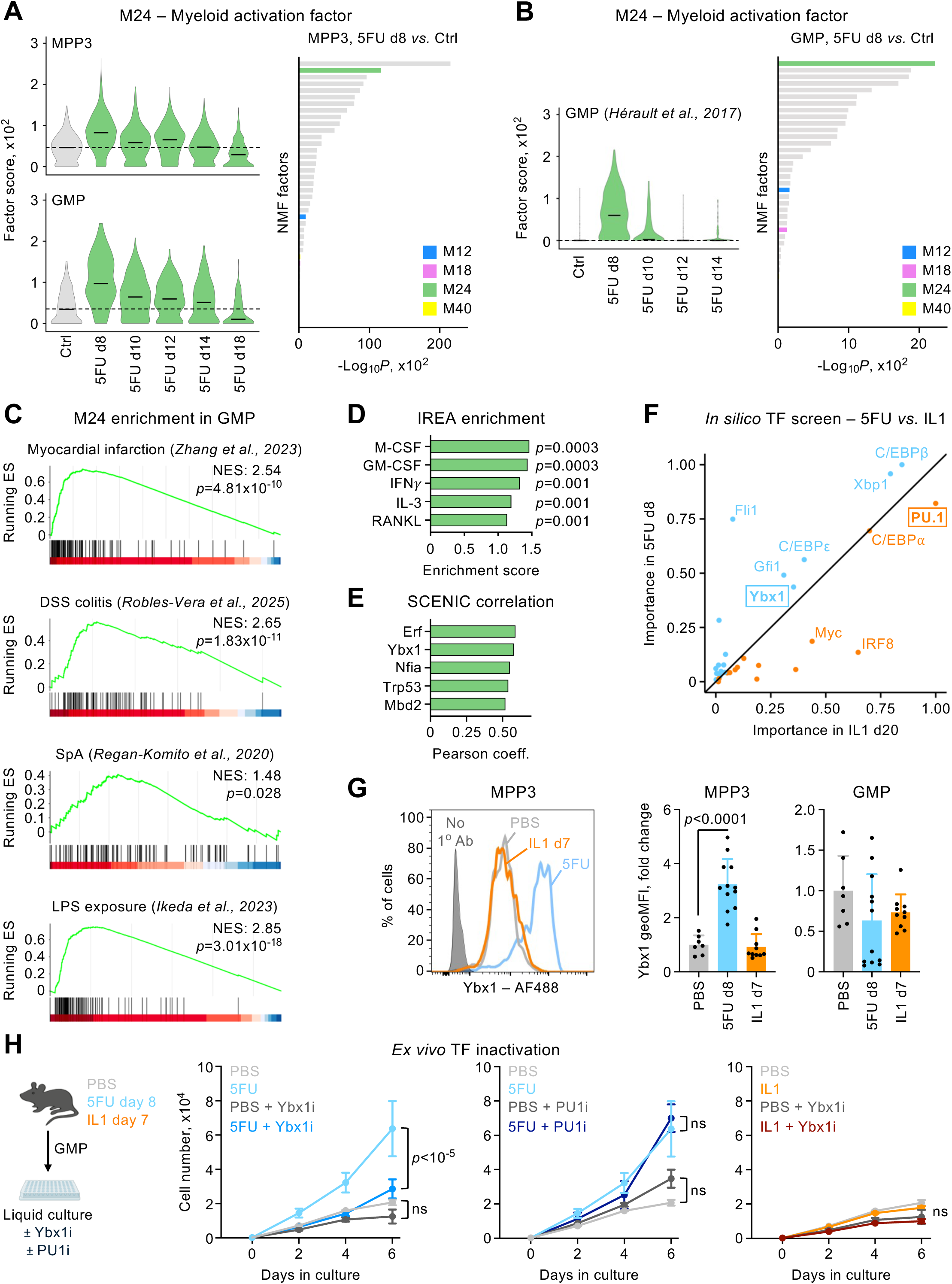
Effector module of myeloid activation is shared across regenerative contexts. (**A**) Violin plots (left) showing NMF M24 cell scores in MPP3 (top) and GMP (bottom) after projection of 10X LSK scRNA-seq data from 5FU treatment timecourse dataset onto NMF model. Bar chart (right) showing *P. values* (Wilcoxon rank sum test with Bonferroni correction) for comparison of factor scores between day 8 and day 0 in MPP3. (**B**) Violin plot (left) showing NMF M24 cell scores in GMP after projection of SMART-seq data from 5FU timecourse dataset onto NMF model. Bar chart (right) showing *P. values* (Wilcoxon rank sum test with Bonferroni correction) for comparison of factor scores between day 8 and day 0 GMP. (**C**) Enrichment plots for top 100 genes loading on M24 in indicated published GMP bulk RNA sequencing datasets. (N)ES: (normalized) enrichment score. *P. values* from permutation testing. (**D**) Bar chart showing results of immune response enrichment analysis (IREA) for M24 genes. *P. values* from two-sided Wilcoxon rank sum test. (**E**) Bar chart showing top 5 Pearson correlation coefficients (coeff.) for NMF M24 scores and SCENIC regulons. (**F**) Scatter plot showing scaled importance of transcription factors (TF) in myeloid differentiation of 5FU or IL-1 treated cells, in a screen of 33 TFs using CellOracle. Values represent the inner product of perturbation score and development flow for each TF, scaled between 0 and 1 for each perturbation. (**G**) Representative flow cytometric image (left) for intracellular staining for Ybx1 in MPP3 treated with either PBS, 5FU at day 8, or IL-1 for 7 days. No 1° indicates control with only secondary antibody staining. Bar charts (right) showing fold change in geometric mean fluorescence intensity (GeoMFI) for Ybx1 compared to PBS in either MPP3 or GMP. Points represent individual mice with mean ± S.D. and *P. values* from one-way ANOVA with *post hoc* Tukey’s test. (**H**) Isolation of GMP from indicated treatment groups, followed by liquid culture with or without small molecular inhibitors of Ybx1 (SU056) or PU.1 (DB2313). Graphs show number of cells in culture over time. Points represent n=4 biological replicates per group with mean ± S.D. and *P. values* from two-way ANOVA with Sidak’s *post hoc* test. See also Figure S12.

### Myeloid activation factor predicts outcome in acute myeloid leukemia

Since M24 was activated in several different inflammatory and regenerative contexts in myeloid progenitors, we asked whether a similar effector module could regulate human hematopoiesis. To establish whether common signatures were activated in human HSPCs exposed to EM perturbations, we used two complementary approaches. First, we created ∼50-gene signatures by identifying top loading genes of our 4 EM-associated mouse NMF factors (M12, M18, M24, M40) that were significantly upregulated in relevant perturbations (mSig) before deriving human orthologs (hSig) (**Fig. 7A; Table S15**). Then, we also searched for common modules in human EM contexts by integrating published BM scRNA-seq datasets from healthy volunteers exposed to G-CSF for 5 days^68^ and BM cells treated *ex vivo* with either LPS or the TLR1/2 agonist Pam3CSK4 for 4 days^69^ (**Fig. 7A**; **Fig. S12C; Table S16**), before mapping to a reference embedding^70^ and performing NMF with 30 factors (**Table S17**). Remarkably, we identified a human factor H2 that was activated in GMP, but not HSC/MPP, by both LPS and Pam3CSK4 (**Fig. 7B**), which shared many key genes with the murine-derived mSig24 (**Fig. 7C**) and in which YBX1 was also identified as a major predicted regulator (**Fig. S12D**). Together, this suggests that the molecular process represented by H2/mSig24 is conserved across mammal species, validating our approach to identify fundamental EM signatures in murine datasets.

**Figure 7.**
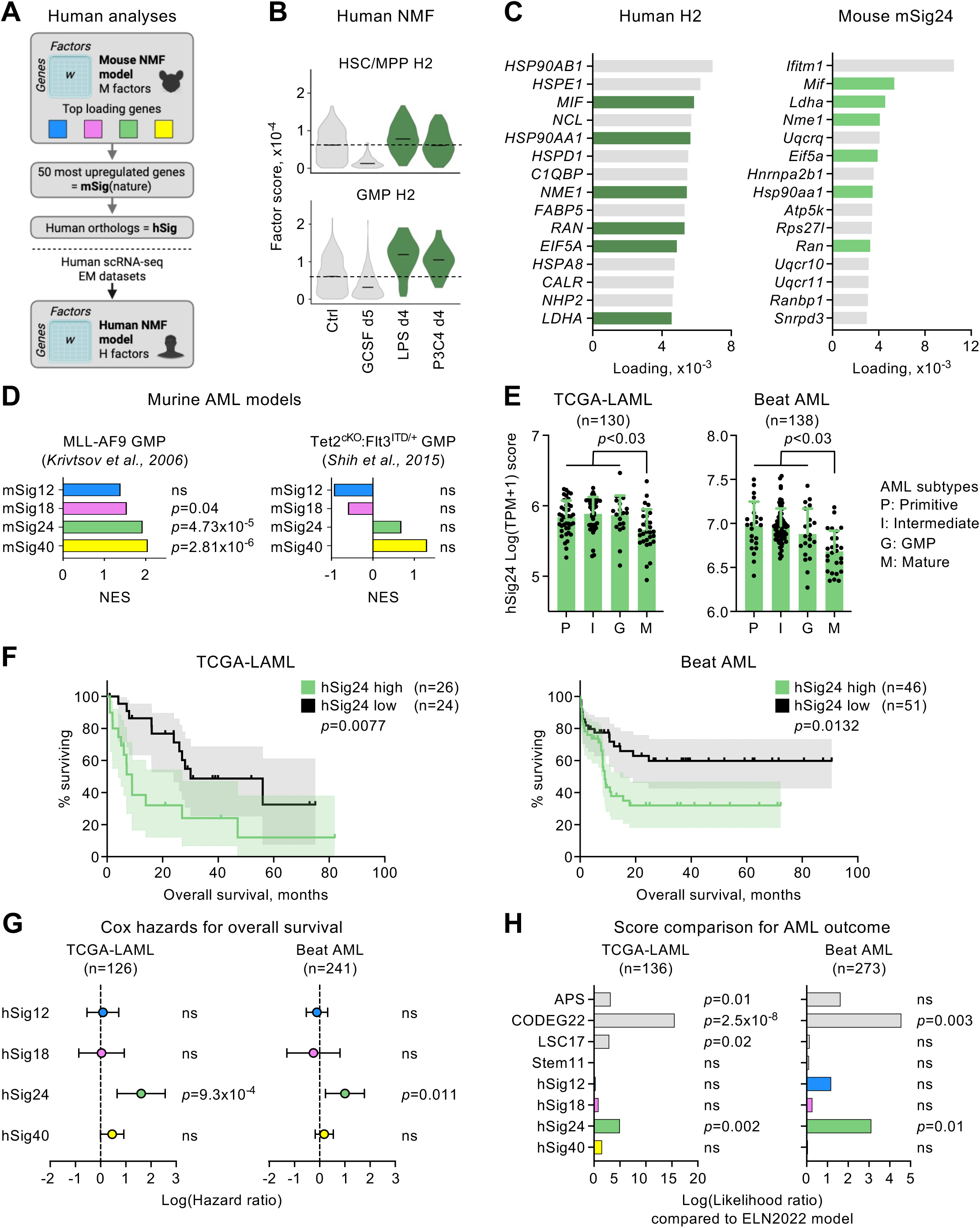
Myeloid activation factor is conserved in humans and predicts outcome in acute myeloid leukemia. (**A**) Outline for generation of murine (mSig) and human EM signatures (hSig) of ∼50 genes from top loading and most differentially regulated genes in NMF model, and for generation of human NMF model of EM perturbations. (**B**) Violin plots showing activity of human NMF factor (H)2 in HSC/MPP (top) and GMP (bottom) in indicated perturbation conditions. (**C**) Bar plots showing top 15 genes by NMF loading coefficient for human H2 and murine signature (mSig)24, with colored bars indicating shared genes. (**D**) Bar charts showing normalized enrichment score (NES) for murine EM signatures in bulk RNA sequencing of GMP from indicated acute myeloid leukemia (AML) models. *P. values* from permutation testing. (**E**) Bar charts showing scores derived by multiplying loading scores for genes in EM signatures with log transcripts per million (TPM)+1 in bulk RNA sequencing samples from TCGA-LAML and Beat AML cohorts. Points show values for individual patients with mean ± S.D. and *P. values* from one-way ANOVA with Tukey’s *post hoc* test. (**F**) Kaplan-Meier curves for overall survival for individuals without TP53 mutations with top and bottom 20% of hSig24 scores in TCGA-LAML and Beat AML cohorts. *P. values* from log rank tests. (**G**) Forest plots showing model coefficients for Cox proportional hazards models for overall survival in TP53 wild type patients in TCGA-LAML and Beat AML cohorts, including scores for EM signatures as covariates alongside age, sex, and previous malignancy or treatment. *P. values* from Cox regression. (**H**) Bar charts showing likelihood ratios for predictive capacity of prognostic models incorporating indicated molecular scores, compared to baseline European Leukemia Network (ELN) 2022 risk scores. *P.* values from ANOVA test for Cox model fits based on the log partial likelihood. See also Figures S12, S13, and S14.

Since SARS-CoV-2 (COVID-19) has been associated with aberrant EM responses, we next tested our signatures among genes upregulated in peripheral blood HSPCs of COVID-19 patients^71^, finding significant enrichment for hSig24 and hSig40, whereas hSig12 was downregulated (**Fig. S12E,F**). Interestingly, hSig24 enrichment was driven particularly by genes related to metabolic activity and stress response, such as *HSPD1*, *NME1*, and *LDHA* (**Fig. S12F**), and not primarily by IFN-response genes described previously in severe COVID-19 cases^72^. Appropriation of molecular processes that drive activation and proliferation is a common strategy of neoplasia, and we next investigated whether our EM signatures were enriched in AML in both mice and humans. First, we found that mSig24 was significantly enriched in an AML mouse model produced by transformation of GMP with the MLL-AF9 fusion protein^73^, but not in the *Tet2^f/f^:Vav1-iCre^+/-^:Flt3^ITD/+^* AML mouse model that originates from more primitive transformed HSPCs^74^ (**Fig. 7D; Table S13**). Using a classification system based on predicted cell of origin^75^, we found that human patients in the TCGA-LAML^76^ and Beat AML^77,78^ cohorts had higher hSig24 scores in the ‘primitive’, ‘intermediate’, and ‘GMP’ groups compared to the ‘mature’ subtype (**Fig. 7E**), consistent with the relative enrichment of M24 in murine GMP and MPP3. Next, we asked whether hSig24 enrichment might predict survival outcomes in AML patients using univariable analysis of different molecular alterations alongside hSig24. We found that both TP53 mutation status (z=4.42, q<0.001, Cox regression with Benjamini-Hochberg correction) and hSig24 (z=3.31, q=0.008, Cox regression with Benjamini-Hochberg correction) were significantly associated with worse overall survival after false discovery rate correction, and that patients with TP53 mutations in the Beat AML cohort had higher hSig24 scores (**Fig. S13A-C; Table S18**). Since TP53 mutations are already known to be associated with worse prognosis in AML^76,79^, we performed two analyses to assess the independence of the contributions of hSig24 and TP53 mutation status. First, we constructed a multivariable Cox regression model in the Beat AML cohort, using the 5 features that were significantly associated with overall survival in the univariable analysis (hSig24, TP53, BCOR, CEBPA, and U2AF1), and finding that hSig24 remained a significant independent predictor of overall survival (**Fig. S13C**). Second, in order to examine the prognostic value of hSig24 independent of TP53 mutation status, we removed all mutant TP53 samples and then compared the overall survival of wild type (WT) TP53 patients with the highest (top 20%) and lowest (bottom 20%) hSig24 scores in the TCGA-LAML and Beat AML cohorts. We found that patients with the highest hSig24 scores had significantly poorer survival in both cohorts (**Fig. 7F**). To evaluate our EM signatures as predictors of overall survival in AML patients, we constructed Cox proportional hazards models that accounted for age, sex, previous malignancy, and treatment history as covariates (**Fig. 7G**) and discovered hSig24 was associated with poorer survival among WT TP53 patients in both TCGA-LAML and Beat AML cohorts, whereas other EM scores did not predict outcome (**Fig. 7G**). We also found that both hSig24 and hSig40 predicted outcome in the pediatric TARGET-AML cohort (**Fig. S14A**), despite the major differences in molecular and clinical features observed between adult and pediatric AML^80^. In contrast, hSig24 was not useful in predicting outcome in lymphoid hematopoietic cancers (**Fig. S14B**) or in most solid cancers, though it was associated with overall survival in patients with lung adenocarcinoma in the TCGA-LUAD cohort^81^ (**Fig. S14C**), in which others have also reported myeloid signatures that inform outcome^82,83^. Finally, by comparing likelihood ratios, we found that hSig24 significantly increased prognostic accuracy over the established ELN2022 clinical score^84^, which is based on mutation and cytogenetic profiles and classifies TP53-mutated AML as a separate disease entity. In fact, hSig24 outperformed comparable gene scores generated through different approaches^85^, such as the APS^86^, LSC17^[87]^, and Stem11^[88]^ scores (**Fig. 7H**). Collectively, these data show that our discovery of a conserved effector module of myeloid activation has direct clinical relevance owing to its ability to predict outcome beyond known factors like TP53 mutation in both pediatric and adult AML.

## DISCUSSION

Here, we exploit a large collection of EM datasets to develop a new approach for HSPC annotation in scRNA-seq datasets (HemaScribe) and to advance a refined quantitative model for hematopoietic differentiation (HemaScape). Our data show great agreement between cell types defined by flow cytometry and molecular profiling as identified by HemaScribe, but imperfect correspondence with density clusters representing stable cell states in transcriptional space as modeled by HemaScape, which mostly incorporate mixed groups of MPPs and lineage-committed progenitors. These results lend support to the emerging notion that many discrete HSPC populations defined by flow cytometry and surface marker expression represent in fact an agglomerate of cells with mixed lineage potential and differentiation stage^89^. One approach to resolve this heterogeneity has been to identify new surface markers that can be used to further subdivide flow-defined populations, as shown for common myeloid progenitors (CMP)^90^. Here, we demonstrate instead that consistent annotation of HSPC cell types across datasets using our toolkit is a powerful new method to resolve this heterogeneity by combining both experimental data and molecular analyses within the same topological structure. This allowed us to develop models explaining how different EM perturbations enhance myeloid output, uncover new shared EM regulatory modules in both mouse and human HSPC hierarchies, and provide a roadmap for investigating a broad range of perturbations in the hematopoietic system.

Using a collection of molecular methods, we show that pathogen-associated EM stimuli like LPS and pIC cause widespread effects on the hematopoietic system, including activation of early HSC/MPP and emergence of novel differentiation pathways, which we validate experimentally. Conversely, the regenerative cytokine G-CSF exerts a focused effect on myeloid-restricted progenitors using only existing differentiation links. These differences are probably explained by multiple factors, including expression of receptors for key mediators of both types of perturbation on different HSPC populations, as well as activation of a more diverse range of downstream molecular processes, as unveiled by our NMF analysis. Functionally, these differences probably also reflect the evolutionary importance attached to severe bacterial and viral infections, for which there is increasing evidence that the nature and extent of HSPC responses can dictate clinical outcome^91^, whereas G-CSF is considered to have a primary role in feedback responses governing homeostatic myeloid cell production^92^.

Through integrative analysis of multiple perturbation conditions, we identify common molecular processes recruited in non-overlapping combinations in different EM responses, suggesting that the hematopoietic system has only a limited number of ways to respond to diverse stimuli. This situation represents an efficient means to encode activation responses in the genome, which leads us to make the following predictions about these shared “EM effector modules”. First, we speculate that key genes and regulators will be subject to strong positive selection as a reflection of their central importance in hematopoietic responses, similar to what is observed for critical immune regulators^93^. Second, the shared nature of molecular processes across EM conditions suggests that they might represent novel activation states, in which cells establish alternative but stable transcriptional networks that deliver demand-adapted effector functions. This is supported by studies showing that termination of hematopoietic activation often requires recruitment of active “cellular braking” mechanisms and not just the removal of the initiating insult^40,94^. Finally, we predict that inflammatory and infectious diseases associated with maladaptive HSPC activation might be sustained or exacerbated by imbalances in the relative activity of different modules and not just by altered activity of single TFs or regulators. Interestingly, many of the molecular processes we describe have received little functional investigation to date. For instance, while Runx1 has been studied for decades in the context of hematological cancers and emergence of embryonic HSCs^95^, its role in EM responses suggested by our evolutionarily conserved M18 factor is currently understudied. Others have also speculated that AP-1 factors might represent common drivers of epigenetic memory after inflammation^96^, suggesting that our shared M12 factor observed in conditions of chronic inflammation could be a driver of central trained immunity. We also identify M24 as a shared activation module among myeloid progenitors, which is an attractive topic for further study as a possible therapeutic target in diseases characterized by maladaptive EM responses. Importantly, we also find that this myeloid activation factor predicts outcome in human adult and pediatric AML, suggesting the molecular process represented by this signature is hijacked in cancer to produce more severe disease manifestations. Interestingly, Ybx1, which we identify as a candidate driver of the myeloid activation factor, is an important dependency in AML cell lines that drives cell proliferation^97^, and the small molecule Ybx1 inhibitor SU056 shows promise for its treatment in initial evaluations^98^. Here, we find that Ybx1 could have important physiological functions in myeloid cell expansion, which is consistent with its known roles in stabilizing transcripts of TFs that drive myelopoiesis, particularly Myc^97,99^. Together, this suggests an evolutionary trade-off in maintaining molecular mechanisms that promote regeneration, but which may be exploited in mutated cancer cells and maladaptive conditions to permit uncontrolled proliferation. Collectively, our work illuminates fundamental regulatory mechanisms in hematopoietic regeneration and identifies a common EM effector module that informs outcome in human disease contexts.

### Limitations of the study

We identify several modules in hematopoiesis that are repeatable across datasets, but, aside from the myeloid activation factor M24, we currently lack information on the functional role and importance of these molecular processes in both steady state and perturbation conditions. We envisage that this will form the basis for follow-up studies using functional genomic approaches to explore the importance of these factors in disease. Although we validate our HemaScribe annotation approach extensively against other resources, we do not possess ground truth data for cell types in most perturbation conditions except for day 8 5FU, meaning that some cell type annotations could be less stable under perturbed conditions. Finally, although our HemaScape tree captures many consensus elements of hematopoietic topology, the cross-sectional nature of our molecular profiling does not provide additional insight on the ongoing debate about the extent of genuine multipotency in the hematopoietic system, which can only be resolved by data modalities that incorporate combined molecular and functional readouts in the same single cells.

## RESOURCE AVAILABILITY

All newly generated datasets supporting this study have been deposited in the Gene Expression Omnibus (GEO) under accession numbers GSE296404 and GSE298048. Previously published datasets analyzed in this study are indicated in the Key Resources Table. An interactive website for exploration of our reference dataset of 10 oligo-hashed flow-isolated HSPC populations is available online at http://3.233.5.190:8050. Functions for HemaScribe annotation of cells in scRNA-seq datasets and mapping to the HemaScape differentiation landscape are available in the R package “HemaScribe” with supporting reference data, available for download at <u>github.com/RabadanLab/HemaScribe</u>. All other analyses were conducted with publicly available software packages, as outlined in the Key Resources Table.

## ACKNOWLEDGEMENTS

We thank Dr. D. Landau and Dr. F. Izzo (New York Genome Center) for providing the original annotation of their dataset; Drs. J. Dick and A. Zeng (University of Toronto) for providing their BM map and classification based on cell deconvolution for human AML samples; Dr. A. Wilkinson (University of Oxford) for providing raw data from CRISPR/Cas9 screens; and Drs. C. Lachowiez and N. Long (Oregon Health and Science University) as well as Drs. Z. Sachs, L. Baughn, and Y. Lee (University of Minnesota) for their assistance with ELN scores for the TCGA-LAML and Beat AML cohorts. We thank M. Kissner for management of the CSCI Flow Cytometry Core facilities, and all members of the Passegué and Rabadan laboratories for critical insights and suggestions. The results shown here are in part based upon data generated by the TCGA Research Network (www.cancer.gov/tcga) and by the Therapeutically Applicable Research to Generate Effective Treatments (www.cancer.gov/ccg/research/genome-sequencing/target) initiative. J.W.S. was supported by EMBO postdoctoral fellowship ALTF-2021-196 and Damon Runyon Cancer Research Foundation DRG-2493-23 (William Raveis Family Fellowship). This work was funded by NIH R35HL135763, R35HL171521, and R01CA255342 to E.P., NIH R35CA253126 to R.R., NIH P01CA285250 to E.P. and R.R., and was supported in part through the NIH/NCI Cancer Center Support Grant P30CA013696 to CUIMC.

## AUTHOR CONTRIBUTIONS

Conceptualization, J.W.S., J.H.F., Z.C., R.R. and E.P.; methodology, J.W.S., J.H.F., Z.C.; investigation, J.W.S., J.H.F., Z.C., O.C.O., A.C., M.A.P., R.Z.; visualization, J.W.S., J.H.F., Z.C., T.L.; funding acquisition, R.R., E.P.; project administration, R.R., E.P.; supervision, R.R., E.P.; writing – original draft, J.W.S., J.H.F., Z.C., R.R., E.P.; writing – review & editing, J.W.S., J.H.F., Z.C., R.R., E.P.

## DECLARATION OF INTERESTS

R.R. is a founder of Genotwin and a member of the SAB of Diatech Pharmacogenetics and Flahy. None of these activities are related to the work described in this manuscript. The other authors declare no competing interests.

## STAR METHODS

### Resource Availability

#### Lead Contact

Further information and requests for resources and reagents should be directed to and will be fulfilled by the Lead Contacts.

#### Materials Availability

This study did not generate new unique reagents.

#### Data and code availability

Single cell RNA sequencing datasets generated in this study have been deposited at GEO. Accession numbers are listed in the Key Resources Table, along with details of all published datasets re-analyzed in this study. An interactive website for exploration of our reference dataset of 10 oligo-hashed flow-isolated HSPC populations is available online at http://3.233.5.190:8050. Functions for HemaScribe annotation of cells in scRNA-seq datasets and mapping to the HemaScape differentiation landscape are available in the R package “HemaScribe” with supporting reference data, available for download at <u>github.com/RabadanLab/HemaScribe</u>. All other analyses were conducted with publicly available software packages, as outlined in the Key Resources Table. Any additional information required to re-analyze the data reported in this paper is available from the lead contacts upon request.

### Experimental Model and Study Participant Details

#### Animals

All animal experiments were conducted at the Columbia University Irving Medical Center (CUIMC) in accordance with approved Institutional Animal Care and Use Committee and in compliance with all relevant ethical regulations. Wild type (WT) C57BL/6J (CD45.2^+^) and B6.SJL-*Ptprc^a^Pepc^b^*/BoyJ (CD45.1^+^, BoyJ) mice were purchased from the Jackson Laboratory and bred in house, like the β-actin-GFP mice^49^. Tg(Gt(ROSA)26Sor-Fucci2 mice^36^ were kindly provided by Dr. H.L. Grimes (Cincinnati Children’s Hospital). Aged WT C57BL/6 mice were obtained from the National Institute on Aging (NIA) when they were 18 months old and used for experiments when they were 24 months old. Unless otherwise stated, mice were 8 to 12 weeks of age when used for experiments. No specific randomization or blinding protocol was used with respect to the identity of experimental animals, and both male and female animals were used in all experiments. Mice were euthanized by CO_2_ asphyxiation followed by cervical dislocation.

### Method Details

#### In vivo assays

5-fluorouracil (5FU) (Sigma Aldrich F6627) was dissolved in sterile PBS (10 mg/ml solution) and administered once at 150 mg/kg by intraperitoneal (i.p.) injection; control mice were injected with PBS. Recombinant human granulocyte colony stimulating factor (G-CSF) (Neupogen, Amgen) was diluted in sterile PBS (50 μg/ml solution) and injected i.p. daily at 5 μg per mouse; control mice were injected with PBS. Recombinant murine interleukin (IL)-1β (PeproTech) was reconstituted in PBS containing 0.1% bovine serum albumin (BSA) (5 μg/ml solution) and injected i.p daily at 0.5 μg per mouse; control mice were injected with PBS/0.1% BSA. Polyinosinic:polycytidylic acid (pIC) (Cytiva) was dissolved in PBS (1.25 mg/ml solution) and injected i.p. at 10 mg/kg every 48 hours; control mice were injected with PBS. Aliquots of pIC were warmed to 65°C for 15 minutes before injection. Recombinant murine erythropoietin (EPO, PeproTech) was injected at 4 U/g i.p. for 2 consecutive days, as described^41^.

#### Complete blood cell counts

Blood was collected by cardiac puncture after euthanasia and immediately transferred into EDTA-coated tubes (Greiner Bio-One, 0.5 ml). Complete blood cell counts were performed with an Oxford Science Genesis analyzer.

#### Flow cytometry of hematopoietic cells

BM cells were obtained by crushing 8 hindlimb, forelimb, and pelvic bones in staining medium composed of Hanks’ buffered saline solution without calcium or magnesium (HBSS, Gibco, 14175079) containing 2% heat-inactivated fetal bovine serum (FBS, Gibco HI FBS, 16140-071). RBCs were removed by lysis with ACK buffer (150 mM NH_4_Cl and 10 mM KHCO_3_), and single-cell suspensions of BM cells were purified on a Ficoll gradient (Histopaque 1119, Sigma-Aldrich). For HSC and progenitor isolation, BM cells were pre-enriched for c-Kit^+^ cells using c-Kit microbeads (Miltenyi Biotec, 130-091-224) and an AutoMACS cell separator (Miltenyi Biotec). Unfractionated or c-Kit-enriched BM cells were then incubated with the following antibodies: CD3e-PE/Cy5 (Invitrogen, 15-0031-63; 1:100), CD4-PE/Cy5 (Invitrogen, 15-0041-81; 1:1600), CD5-PE/Cy5 (BioLegend, 100610; 1:800), CD8a-PE/Cy5 (Invitrogen, 15-0081-81; 1:800), CD11b-PE/Cy5 (Invitrogen, 15-0112-81;1:1600), CD19-PE/Cy5 (BioLegend, 115510; 1:400), B220-PE/Cy5 (Invitrogen, 15-0452-81; 1:800), Gr1-PE/Cy5 (Invitrogen, 15-5931-82, 1:800), Ter119-PE/Cy5 (Invitrogen, 15-5921-83; 1:400), c-Kit-APC/Cy7 (BioLegend, 105826; 1:800); Sca-1-BV421 (BioLegend, 108128; 1:400); CD150-BV650 (BioLegend, 115931; 1:200); CD48-AF700 (BioLegend, 103426; 1:400); Flt3-PE (Invitrogen, 12-1351-82; 1:100) or Flt3-biotin (Invitrogen, 13-1351-85; 1:100) followed by streptavidin (SA)-APC (BioLegend, 405207; 1:200), CD127-PE (Invitrogen, 12-1271-82), CD34-FITC (Invitrogen, 11-0341-85; 1:50) or CD34-biotin (BioLegend, 119304; 1:100), CD16/32-PE/Cy7 (BioLegend, 101317; 1:800), CD41-BV510 (BioLegend, 133923; 1:200), and CD105-BV786 (BD, 564746; 1:200). For analysis of apoptosis, cells were washed after initial staining then incubated with Apotracker Green (BioLegend, 427401) according to the manufacturer’s instructions. For peripheral blood chimerism analyses after MPP4 transplant, blood was collected by retro-orbital sampling under isoflurane anesthesia, and RBCs were lysed with ACK buffer. Cells were then stained with CD11b-PE/Cy7 (Invitrogen, 25-0112-82; 1:800), Gr1-e450 (Invitrogen, 57-5931-82; 1:400), B220-APC/Cy7 (Invitrogen, 47-0452-82; 1:800), CD3-APC (Invitrogen, 17-0032-82; 1:100) and Ter119-PE/Cy5 (Invitrogen, 15-5921-83; 1:400) together with CD45.2-FITC (Invitrogen, 11-0454-85, 1:400) and CD45.1-PE (Invitrogen, 12-0453-83, 1:400). For peripheral blood chimerism analysis after GFP^+^ HSC transplant, blood was collected by retro-orbital sampling into 1.5 ml tubes containing 20 μl of 0.5M EDTA solution, pH 8 (Invitrogen 15575-038). Tubes were centrifuged at 250 *g* for 10 mins, after which the platelet-rich plasma was removed into a new tube and stained with Ter119-PE/Cy5 (Invitrogen, 15-5921-83; 1:400), CD61-PE (BioLegend, 104308, 1:400), CD150-APC (BioLegend, 115910, 1:400), with acquisition also in the GFP channel. Before analysis, stained cells were resuspended in staining medium containing 1 µg/ml propidium iodide (Sigma, P4170) for dead cell exclusion. Isolation of specific cell types was performed on a FACS Aria II SORP (BD) using double sorting for purity. Flow cytometric analyses were performed on a NovoCyte Quanteon or NovoCyte Penteon cell analyzer (Agilent). Data collection was performed using FACSDiva (v9) or NovoExpress (v1.6.2), and analysis was conducted using FlowJo (v9/v10).

#### Flow cytometry for Ybx1

For intracellular staining of Ybx1, c-Kit-enriched BM cells were stained for surface markers for 30 minutes at 4°C (CD150-APC [BioLegend 115909, 1:400], CD48-AF700 [BioLegend, 103426; 1:400], c-Kit-APC/Cy7 [BioLegend, 105826; 1:800], Sca-1-Pacific blue [BioLegend 108120, 1:200], Flt3-PE [Invitrogen 12-1351-82, 1:100], CD16/32-PE/Cy7 [BioLegend, 101317; 1:800], and lineage stains in PE/Cy5 as described above) then fixed and permeabilized using the Foxp3/Transcription Factor Fixation/Permeabilization and 10X Permeabilization solutions (ThermoFisher Scientific, 00-5523-00) according to the manufacturer’s instructions. After this, cells were stained with rabbit anti-mouse Ybx1 antibody (Abcam, ab76149, 1:200 dilution in 1X Permeabilization solution) for 1 hour at 4°C. After washing, cells were stained with goat anti-rabbit IgG AF488 (Invitrogen, A11008, 1:400 dilution) for 30 minutes at 4°C. Cells were then washed and re-suspended in 1X Permeabilization buffer for acquisition on a NovoCyte Penteon (Agilent).

#### Liquid culture of GMPs

GMPs were first sorted into 1.5 ml tubes containing HBSS with 2% FBS using a FACS Aria II (BD) on “yield” setting and then re-sorted on “4-way purity” to deposit 250 cells per well of a 96 well U bottom plate filled with 200 μl of Iscove’s modified Dulbecco medium (IMDM, Invitrogen 31-980-097) containing 5% FBS, 100 U/ml penicillin and 100 μg/ml streptomycin (Thermo Scientific, 15140122), 0.1 mM non-essential amino acids (Fisher Scientific, 11-140-050), 1 mM sodium pyruvate (Fisher Scientific, 11-360-070), 2 mM L-glutamine (Fisher Scientific 35-050-061), 50 μM 2-mercaptoethanol (Sigma, M7522), and the following cytokines (all from PeproTech): IL-3 (10 ng/ml), granulocyte–macrophage colony-stimulating factor (GM-CSF; 10 ng/ml), stem cell factor (SCF; 25 ng/ml), IL-11 (25 ng/ml), Flt3L (25 ng/ml), thrombopoietin (TPO; 25 ng/ml), and erythropoietin (EPO; 4 U/ml). In some wells, the Ybx1 inhibitor SU056 (Selleck Chemicals, E1331) was added at 500 nM, or the PU.1 inhibitor (Glixx Labs, 2170606-74-1) at 50 nM. Cells were counted every 2 days by removing 100 μl of medium and mixing with 50 μl of HBSS containing 2% FBS and 1 μg/ml propidium iodide before counting on a NovoCyte Penteon using the “absolute count” setting.

#### Transplantation

All transplant recipients were congenic CD45.1^+^ BoyJ mice that were either lethally (5.25 Gy x2) or sublethally (4 Gy x2) irradiated before transplantation using a MultiRad225 irradiator (Precision X-Ray Irradiation). To evaluate platelet output, 500 HSCs isolated from β-actin-GFP mice were injected per lethally irradiated BoyJ recipient by retro-orbital injection under isoflurane anesthesia alongside 600,000 Sca-1-depleted BoyJ helper BM cells. To evaluate MPP4 output, 5,000 MPP4s isolated from WT C57BL/6 mice were injected per sublethally irradiated BoyJ recipient by retro-orbital injection under isoflurane anesthesia alongside 600,000 Sca-1-depleted BoyJ helper BM cells. Starting the day after transplant, some recipients were injected with pIC (2 injections, 48 hours apart) or G-CSF (4 injections, every 24 hours). For MPP4 transplants, a second cycle of injections was conducted starting on day 21 after transplant. Blood samples were obtained by retro-orbital bleeding starting 7 days after transplant, and then every 4-5 days thereafter. Transplant recipients received antibiotic water (polymyxin B [Sigma Aldrich P4932] 1000 U/ml and neomycin [Sigma Aldrich N1876] 1.2 mg/ml) for 4 weeks after transplant.

#### Generation of scRNA-seq datasets

30,000 to 50,000 LK or LSK cells were sorted into 1.5 ml tubes containing a 1:1 mixture of HBSS and FBS and then rested for 1 hour on ice before pelleting at 350 x *g* for 5 minutes at 4°C. The supernatant was removed down to a volume of 40 μl before GEM generation and 3’ RNA library preparation were performed according to 10X Genomics protocol CG000315 Rev E, targeting 5000 cell data recovery. RNA libraries were pooled 1:1:1:etc., sequenced on an Illumina NovaSeq 6000 (2×150 bp, 2.5 billion reads), and aligned using Cell Ranger (v.7.0.1) to mouse genome mm10. Library concentrations and fragment sizes were evaluated using Qubit dsDNA HS assay kit (ThermoFisher Scientific) and TapeStation D5000 DNA ScreenTape analysis (Agilent).

#### Quality control for scRNA-seq data

Standard quality control filtering was performed on each newly generated single cell RNA-sequencing dataset, removing empty droplets (cells expressing fewer than 200 unique genes) and low-quality cells based on outliers for total counts detected (in log scale), number of unique genes detected (in log scale), percentage of counts in top 50% of expressed genes, and percentage of mitochondrial counts^100^. Thresholds were automatically set at ±5 median absolute deviations (MADs) for each quality control variable. Additionally, cells with >8% mitochondrial counts were also excluded. The hashing oligo-tagged reference samples were demultiplexed using *MULTI-seq*^101^ as implemented in Seurat v5^102,103^, following which doublets and negative cells were removed and singlet cells were assigned to one of the ten hashing-labelled cell types. For all other non-hashing datasets generated in this study, doublet removal was performed using *scDblFinder*^104^.

#### HemaScribe: murine bone marrow cell type annotations

We developed HemaScribe, a hierarchical strategy to annotate and classify cell types within the mouse bone marrow by integrating multiple well-established methods for dissecting the identities of single cells with bulk and single cell transcriptomic references into one pipeline. HemaScribe consists of three modules. The first is a hematopoietic cell filter based on gene set scores calculated using the *UCell::ScoreSignatures_UCell* function^105^ with a list of hematopoietic marker genes that were selected from genes differentially expressed between all hematopoietic and non-hematopoietic cells in the Tabula muris droplet datasets, manually supplemented with additional genes for erythropoiesis (**Table S3**). Depending on tissue source and contamination a sample may contain non-hematopoietic cells and it is important to remove such cells to improve the efficiency of our subsequent classifiers which are trained solely on hematopoietic cell types. Therefore, after scoring, cells with a hematopoietic score below a manual threshold can be excluded from downstream analysis, and we found that a threshold of ∼0.1 was appropriate for most BM datasets. Cells that pass the hematopoietic cell filter are then classified into broad BM populations using *SingleR*^106^ based on their correlation with a curated set of bulk RNA-sequencing references^16–18^ described in **Table S1**. This was principally done to extract a subpopulation of bone marrow cells corresponding to the early progenitors that align with the single cell LK and LSK reference datasets in the next step but is also useful for resolving other cell type populations (e.g., GMP and GP) in the marrow. Lastly, cells that are identified as “HSPCs” by the broad-level classifier are assigned a finer cell type using Seurat’s integrative analysis based on anchor integration^107^ with our newly generated WT single cell RNA-sequencing datasets with oligo-hashed labels. These reference datasets were created by separately concatenating the WT and 5FU-treated samples and processed using the Seurat functions *NormalizeData*, *FindVariableFeatures*, *FindIntegrationAnchors*, and *IntegrateData* with 3000 variable features and 3000 anchor features each. The integration step was performed with the Seurat commands *FindTransferAnchors* and *TransferData* using 30 principal components. After integrating our query dataset with the reference, the oligo-hashed labels are transferred onto each cell in the query. For ease of interpretation, we grouped together overlapping cell type labels in the broad-level and fine-level annotations to obtain a final combined annotation. We also developed a variant of HemaScribe using the 5FU reference hashing label datasets instead of the WT datasets.

To assess the performance of the fine classifiers, we computed the per-cell type precision and recall on the training datasets as well as on held-out 5FU datasets. We also validated our HemaScribe annotations against previously published hematopoiesis datasets, including a recent transcriptomic atlas of mouse Lin^-^ BM cells^25^ sequenced using the Chromium platform from 10X Genomics, a dataset of LSK cells subjected to CITE-seq^24^, and a collection of bulk RNA sequencing samples from purified HSPCs^12,23^. For the CITE-seq data, a value of 1 was added to the antibody-derived tag (ADT) data, which were then normalized using the “CLR” option in the *NormalizeData* function in Seurat, with margin=2. Cutoff limits were applied for Flk2 (CD135), CD48, and CD150 by visual inspection, with the following values: MPP4: CD135>0.6 and CD150<1; MPP3: CD135<0.5 and CD150<1.5 and CD48>2.5; MPP2: CD135<0.5 and CD150>1.5 and CD48>2.5; HSC: CD135<0.5 and CD150>1.5 and CD48<2.5; and ST-HSC: CD135<0.5 and CD150<1.5 and CD48<2.5. To compare the results of HSPC-sorted cells with HemaScribe, we calculated the adjusted Rand index (ARI) for cell assignments using the *adjustedRandIndex* function from the R package *mclust*. To evaluate associations between HemaScribe and bulk RNA sequencing samples, we first obtained pseudobulk profiles for each HemaScribe population from a control dataset incorporating 3 steady state WT LK and 3 LSK datasets using the *AggregrateExpression* function in Seurat, and we then calculated correlations between these profiles and bulk RNA sequencing samples using the *clustify* function from the R package *ClustifyR*, using the top 2,000 highly variable genes from the scRNA-seq dataset. Next, we scaled each bulk RNA sample and identified its greatest correlate among the HemaScribe populations, and we used these results to calculate the overall ARI for cell types represented in both datasets. In addition to testing the accuracy of our annotations on these external benchmarks, we also examined the internal consistency of our annotations by adapting the cluster stability measures from *scclusteval*^108^. Briefly, we repeatedly sub-sampled 80% of our control datasets 100 times, re-ran HemaScribe on each sub-sample, and recorded the cell type annotations from each run. Next, we computed the Jaccard similarity for each imputed subgroup in the subsampled dataset with the corresponding subpopulation imputed from the full dataset. A high median Jaccard similarity across all the runs for a given cell type means that the annotations for that cell type are robust and stable. We also computed the Jaccard similarity for each subgroup with the off-target subpopulations in the full dataset. A low median Jaccard similarity here means that discordant cell type assignments are uncommon. Finally, we ran HemaScribe on a variety of normal and perturbed hematopoiesis datasets (**Table S2**). After filtering for low quality cells, the datasets generated in this study were preprocessed using the Seurat commands *NormalizeData* and *FindVariableFeatures* with default parameters. The previously published HSPC CITE-seq dataset^24^, EPO dataset^41^, Tabula muris datasets^26^, mouse Lin^-^ BM dataset^25^, and BM stroma dataset^27^ were re-preprocessed using the same methods, except that the list of included cells were retrieved from the original analyses. For all other external datasets, we used the original pre-processed data from the authors.

#### Evaluation of cell frequency changes

To evaluate the statistical significance of changes in HSPC frequency annotated by HemaScribe in perturbation and matched control datasets, we used functions from the R package *scDC*, as outlined by the authors^43^. First, we performed differential composition analysis using *scDC_noClustering* with 500 bootstraps, using the fine annotation from HemaScribe, in which we subdivided GMPs into constituent mGMP, GP, and cMoP. Next, we used *fitGLM* to fit generalized linear models with the CLP cell type as the reference category, and we took p-values from the fixed effect pooled results.

#### HemaScape: Trajectory analysis of hematopoiesis using DensityPath

To investigate the mechanisms underlying hematopoiesis, we employed DensityPath^29^ to reconstruct the optimal cell state-transition trajectory. This method identified a geodesic minimum spanning tree (MST) of representative cell states (RCS) on the density landscape, establishing a least-action path characterized by minimal transition energy associated with cell fate decisions. Prior to applying DensityPath, we integrated four hashing oligo-tagged reference samples using Seurat v5. Principal component analysis (PCA) was performed on the integrated dataset, and dimensions were selected by identifying the largest eigenvalue 𝜆*_i_* where the difference between consecutive eigenvalues fell below a threshold of 5 x 10^-4^. The resulting 25-dimensional PCA space was then used for non-linear dimensionality reduction via Elastic Embedding (EE)^28^, a method that preserves both local and global data structures, to visualize the intrinsic structure of the scRNA-seq data in a 2-dimensional embedded space. In the EE-embedded space, DensityPath estimated the density surface and applied level-set clustering (LSC) to identify distinct high-density cell clusters, designated as RCSs^109^. The cell state-transition trajectory was reconstructed by determining the MST of the RCS peak points, with geodesic distances computed on the density landscape. To resolve regions with low-density structures, such as the basophil lineage, which were obscured within the global structure, we extracted cells mapped to the GMP lineage branch and reapplied DensityPath to capture fine-scale structures. This process identified basophil-specific local clusters, which were then integrated with the global trajectory to yield a refined and comprehensive cell trajectory. Each cell was subsequently mapped to the nearest waypoint along the trajectory based on its geodesic distance. Using the fixed peak point of the HSC cluster as the starting cell, we computed pseudotime by calculating the geodesic distance between the projected positions of the starting cell and any given cell along the state-transition path. While LSC effectively extracted high-density RCSs as landmarks of the complex density landscape, and the identified RCSs provided an effective representation of subpopulations and gene expression, LSC did not fully capture cell type divisions. To address this, we employed mean-shift (MS) clustering^110,111^, a density-based approach, to refine clustering. Instead of clustering cells directly, MS was applied to the waypoints along the trajectory, and cluster labels were then assigned to the corresponding cells mapped to those waypoints. For the basophil lineage, waypoints along the GMP branch were further refined via MS clustering to enhance clustering granularity. The final cell clustering results are referred to as density clusters. After obtaining cell cluster labels for the reference dataset, we used Symphony^112^ to map additional scRNA-seq datasets onto the reference. This approach facilitated the transfer of MS-based density cluster labels and EE embedding to these datasets. Using the predicted EE embedding, cells were mapped onto the cell state-transition path, enabling pseudotime calculation based on their geodesic distances.

In our R package *HemaScribe*, we provide a *HemaScribe* function to annotate cells and a *HemaScape* function to map query scRNA-seq datasets to our EE embedding, identify density clusters, and calculate pseudotime. This package is compatible with Seurat v5 objects and can be found at <u>github.com/RabadanLab/HemaScribe</u> along with the reference data.

#### Validation of trajectory robustness and accuracy through tree similarity analysis

To assess the robustness and accuracy of the HemaScape tree, we compared it with the directed graph reported by Kucinski et al. (2024)^31^ by analyzing the similarity between their cell type distance profiles. For each pair of cell types annotated by HemaScribe, we first identified all graph nodes that contained either of the two cell types, excluding any nodes where the proportion of a given cell type was below 1%. We then computed the shortest path between every pair of nodes, where one node contained cell type 1 and the other contained cell type 2. The overall distance between the two cell types was calculated as the weighted average of these shortest path distances.

Specifically, for a given directed tree or graph 𝑇, the cell type distance from cell type 𝑐𝑡_1_ to cell type 𝑐𝑡_2_ was defined as:

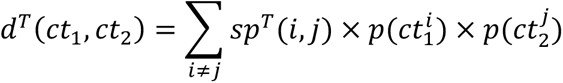

where 𝑐𝑡_1_ and 𝑐𝑡_2_ represent the two cell types being compared, 𝑖 and 𝑗 denote nodes in the tree or graph 𝑇, 𝑠𝑝*^T^*(𝑖, 𝑗) is the shortest path distance from node 𝑖 to node 𝑗 on 𝑇, and 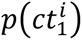and 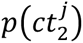 are the proportions of cell types 𝑐𝑡_1_ and 𝑐𝑡_2_ in nodes 𝑖 and 𝑗, respectively. For the Kucinski et al. graph, shortest paths were computed within the network, with edges weighted by the negative exponential of differentiation flux, reflecting transition probabilities. In the HemaScape tree, shortest paths were computed along the tree structure, with edges weighted by geodesic distance, capturing hierarchical relationships. This approach ensures that the calculated distances reflect both the topological structure of the trajectory and the relative abundance of each cell type in the respective nodes.

We evaluated the similarity between the cell type distances derived from the HemaScape tree and those from the Kucinski et al. graph by calculating the correlation between them. To assess statistical significance, we performed 10,000 random permutations of the HemaScape tree. In each permutation, we preserved the start nodes (which are predominantly composed of HSCs) while randomly reassigning connections among the non-start nodes. For each permutation, we recalculated the correlation between the cell type distances of the permuted HemaScape structure and those of the Kucinski et al. graph. The p-value was determined as the proportion of permuted correlations that exceeded the observed correlation, ensuring that the observed similarity was not attributable to random chance. This analysis provided a robust validation of the HemaScape trajectory, confirming its consistency with an established model of cellular differentiation.

#### Comparison of cell label entropy in tree structures

We validated the consistency between our annotations and the clusters found in the tree models obtained from each scRNA-seq dataset by computing the ARI between the node membership for each tree and the fixed HemaScribe annotation using the function *sklearn.metrics.adjusted_rand_score*^113^.

#### PAGA trajectory and connectivity changes with different perturbations

To analyze the connectivity among the MS-based density clusters, we employed Partition-based Graph Abstraction (PAGA)^30^, a graph-based method that abstracts high-dimensional manifolds inherent in scRNA-seq data and quantifies connectivity between density clusters. For hashing oligo-tagged samples, we performed PAGA using the first 50 principal components (PCs) and density clusters after data integration. To align with the DensityPath framework, we visualized the PAGA graph backbone in the embedded EE space, where edge weights represent the PAGA connectivity between cell groups. To assess the effects of different perturbations, we applied PAGA analysis to paired treated-control samples. For each sample, we used the first 50 PCs derived from the scRNA-seq data and density clusters predicted by *Symphony*. Connectivity changes were evaluated by comparing the edge weights between treated and control conditions, enabling us to quantify the perturbation effects on cell differentiation.

#### Evaluation of differentiation tendency

To evaluate transcriptomic bias towards terminal differentiation, we used *Palantir* v1.4.0 as described by the authors^37^. For each dataset, we used the EE embedding created by *Symphony* mapping onto the HemaScape reference dataset, and we defined cells representing start and end points by manual selection; cells used for each dataset are shown in **Table S7**. Possible terminal points were therefore defined in the GP, cMoP, erythroid, megakaryocyte, CLP, and basophil branches. We scaled *Palantir* entropy for cells in each dataset for presentation.

To estimate quantitative changes in hematopoietic differentiation, we created random walk models based on the simplified tree structure outlined above, containing 15 nodes, as described above. For each node, we estimated proliferation rates using a transcriptomic score of cell cycle activity. To do so, we followed the tutorial from the R package *UCell*: we imported count matrices from 6 test datasets (representing 3 WT steady state LK and 3 LSK datasets) into *SingleCellExperiment* objects and scored each cell with the KEGG Cell Cycle geneset (**Table S5**) using *ScoreSignatures_UCell* function. Using our HemaScribe annotation for the same datasets, we then calculated the mean cell cycle score for each cell type. In parallel, we measured the cell cycle status of the same cell types by flow cytometry in Fucci2 mice^36^ that express fluorescent reporters to indicate the cell cycle phase of each cell, identifying actively cycling cells by their expression of Geminin-mVenus. Next, we explored the relationship between cell cycle score and Fucci2 data by fitting linear, logarithmic, and quadratic functions to these data and calculating the Akaike information criterion (AIC) for each. The logarithmic function defined by *y*=0.0172.log(*x*) + 0.0298 had the optimal AIC (data not shown), and we selected this relationship for use with all datasets. For cells assigned to each node in each dataset, we then interpolated the cell cycle activity using the mean cell cycle score for each node using this function. Values greater than 1 (representing 100% of cells engaged in the cell cycle) were assigned a value of 1. To estimate cell death rate in each node, we first estimated cell death rate in each HSPC cell type by flow cytometry using the Apotracker Green dye (BioLegend, 427401) in control mice and in mice subjected to each perturbation that we studied. Next, we calculated an average cell death rate for each node based on its HemaScribe cell type composition and the cell death rates measured by flow cytometry. For this purpose, GP, cMoP, mGMP, and basophils identified by HemaScribe were all treated as GMP, whereas immature B cells were considered as CLP. To estimate the likelihood of cell distribution from an upstream node to its downstream nodes, we used the connectivity matrices generated for each dataset by PAGA analysis, as outlined above. For this purpose, we set all values below the diagonal to 0 (enforcing unidirectional flow), we filtered out connections that did not appear in our tree structure, and we normalized connectivity for each node so that all its downstream connections summed to 1. To estimate differentiation rate for each node, representing the likelihood of cells proceeding from one node to any downstream node(s), we implemented an iterative stochastic simulation approach, where we calculated the size of each node in each dataset using stochastic samples of the following parameters driven by binomial processes: proliferation rate, cell death rate, differentiation rate (set to an arbitrary value of 0.3 in the first iteration), and cell distribution using the normalized connectivity matrix. The starting node size for each iteration was derived from the actual size of each node in LK datasets, which were taken to represent the natural frequency of HSPCs. Cells transitioning from an upstream node were included in the size calculation for their downstream node(s). After each iteration, the error in node size was calculated by comparing final node size with the actual node size, and the differentiation rate was then adjusted according to the direction of the error. Each simulation was run for 10,000 iterations across 10 runs, and differentiation rates were averaged for the final 50 iterations for each run to generate final values for each node. Node parameters used for simulations are provided in **Table S6** (steady state) and **Table S9** (perturbations).

#### Evaluation of transcriptional response

To understand which HemaScape nodes were mounting the greatest transcriptional responses to perturbation, we used the function *calculate_auc* in the R package *Augur*^44^, entering the perturbation as *label_col* and the HemaScape node identity as *cell_type_col*.

#### Inference of transcription factor activity

To estimate transcription factor activity in transcriptomic data, we performed gene regulatory network inference using SCENIC via the fast Python implementation *pySCENIC*^32,114^. Starting with the scRNA-seq expression matrices converted to loom format, we ran the commands *pyscenic grn*, *pyscenic ctx --mask_dropouts*, and *pyscenic aucell* with the auxiliary mm10 and hg38 genome annotations and v10 motif datasets retrieved from https://resources.aertslab.org/cistarget/. To estimate the effects of removing critical transcription factor nodes, we used *CellOracle* v0.18.0^[64]^ implemented via Docker image and using the base gene regulatory network generated by the authors from HSPCs^115^. For each dataset, we used the EE embedding generated by *Symphony* mapping onto the HemaScape reference dataset, as outlined above. We performed in silico perturbations as outlined in the tutorial, and we visualized the results of single perturbations using a digitized grid, for which scale and *min_mass* values are shown for each dataset in **Table S7**. To evaluate the relative importance of different transcription factors in myelopoiesis in different perturbation conditions, we performed an *in silico* screen for a candidate list of 33 transcription factors selected for their known importance in myelopoiesis (**Table S14**). For each dataset, we evaluated the impact of *in silico* perturbations relative to developmental flow defined by calculation of a pseudotime gradient starting in the HSC cluster, with selected starting cells shown in **Table S7**. For the *in silico* screens, we then defined a myelopoiesis trajectory that included the following density clusters: 1, 2, 3, 9, 10, 11, 12. After running the screens, we extracted the negative perturbation score p-values for each transcription factor (representing the significance of disruption relative to developmental flow in the myelopoiesis trajectory) and scaled them between 0 and 1 for each perturbation.

#### Analysis of epigenetic poising

To test whether cells in any HemaScape node showed poised chromatin regions for response to perturbation, we first used the *FindMarkers* function in *Seurat* to derive lists of upregulated genes for all HSPCs together in perturbation datasets compared to their matched controls, excluding any cells labeled as “NotHSPC” in the HemaScribe fine classifier. Next, we used the function *promoterRegions* from the R package *Rsubread* v2.22.1 to derive promoter regions for the murine mm10 genome for the top 500 upregulated genes, which we defined as regions extending 2000 bp upstream and 200 bp downstream from the transcription start site. We also obtained promoter regions for 500 randomly selected genes to act as a measure of background accessibility. Next, we reanalyzed a published single-cell LK/LSK multiome dataset^23^. Briefly, samples for steady state adult LK and LSK cells were analyzed in *Signac* v1.14.0^[116]^ with the following QC parameters: ATAC count >1000 and <100,000 per cell, nucleosome signal <2, TSS enrichment >1. Samples were integrated using the *IntegrateEmbeddings* function, and peaks were called separately for each dataset with MACS2 then combined to create a merged peak file, which was used to count peaks in the *Signac* object. After integration, principal component analysis was performed to identify 30 components, followed by UMAP using components 2 through 30. The RNA assay was processed as described above for scRNA-seq datasets and used to annotate cell types with HemaScribe. Next, we extracted all peaks from the *Signac* object and intersected these regions with the promoter regions for genes upregulated in each perturbation using the *intersect* function in *bedtools* v2.31.1, with the options -wa -a, to retain *Signac* peaks that overlapped with promoter regions. Next, we scored the accessibility of the overlapping promoters in the *Signac* object using the *AddChromatinModule* function in *Signac*. We extracted these scores for each HemaScape node and calculated the mean difference in the accessibility score for each perturbation compared to the sample of promoters from 500 random genes to correct for differences in background accessibility across cell types.

#### Non-negative matrix factorization in mouse and human datasets

To uncover possible shared molecular signatures in emergency myelopoiesis, we first integrated 25 separate scRNA-seq datasets, comprising LK and LSK datasets from 9 different perturbation conditions (**Table S10**). After applying quality control procedures outlined above, these datasets were transformed with *SCTransform* in Seurat v5 and then integrated using the *PrepSCTIntegration*, *FindIntegrationAnchors*, and *IntegrateData* functions. We created a shared list of highly variable genes across all these datasets by also running the *SelectIntegrationFeatures* function with a liberal threshold of 10,000 genes, from which 7,184 genes were retained. From the integrated dataset, we then extracted the SCT-normalized layer that was subsetted against the list of highly variable genes and performed non-negative matrix factorization (NMF) using the *nmf* function from the R package *RcppML* v0.5.6^[117]^. The optimal number of factors was selected by running cross-validation using the *crossValidate* function. The *h* matrix was added to the integrated Seurat object as metadata, whereas the *w* matrix was added as a dimensionality reduction. Each NMF factor was characterized by performing the following procedures: (1) assessment of variance according to HemaScribe cell type and perturbation by calculation of the Kruskal-Wallis test statistic using the *kruskal.test* function from the *stats* R package v4.3.1, (2) evaluation of Gene Ontology biological pathway (GO:BP) enrichment in the genes loading on each factor, using the *GSEA* function from the R package *clusterProfiler* v4.8.3^[118,119]^, with the *scoreType* option as “pos” to account for the absence of negative values, (3) evaluation of cytokine dictionary enrichment by Immune Response Enrichment Analysis (IREA)^55^ (www.immune-dictionary.org) in its interactive interface, using the “macrophage” cell type, and (4) Pearson correlations between *h* matrix scores and SCENIC transcription factor regulon activity for each factor in the same integrated dataset using the *cor* function in *stats*. To understand how NMF factors changed in activity with myeloid differentiation, we also calculated the most active factors based on *h* matrix scores across the pseudotime gradient created with the EE embedding as described above, which were presented as stream plots. To determine the extent of evolutionary conservation of top loading genes in the murine NMF model, we used the *getBM* function in *biomaRt* v2.65.0 to query the Ensembl database of species orthologs for the top 100 loading genes for M12, M18, M24, and M40. We extracted information for *Homo sapiens*, *Danio rerio*, *Xenopus tropicalis*, *Gallus gallus*, *Rattus norvegicus*, *Astyanax mexicanus*, *Poecilia formosa*, *Tetraodon nigroviridis*, *Takifugu rubripes*, *Gadus morhua*, *Pan troglodytes*, *Tursiops truncatus*, *Notamacropus eugenii*, *Podarcis muralis*, *Latimeria chlamunae*, and *Petromyzon marinus*, and we then determined if there was at least 1 annotated ortholog for each of the murine genes. We performed the same analysis with 100 randomly selected genes to represent the background level of conservation and, for presentation purposes, we obtained a dendrogram of evolutionary relationships of vertebrate species from FigShare (https://figshare.com/articles/dataset/Vertebrates_species_trees/22736555). The same procedures were followed for creation of a NMF model using human perturbation datasets, listed in **Table S16**. The *w* matrices from NMF models showing gene loading scores are provided in **Table S11** (mouse) and **Table S17** (human). To project our mouse NMF model onto independent query datasets (listed in **Table S12**), we intersected genes present in the query dataset and NMF model and then performed non-negative least squares regression using the *w* matrix and the query expression matrix to reproduce a pseudo-*h* matrix for the query dataset. This was implemented with the *nnls* function in the R package *nnls* (v1.6). Scores for different perturbations and cell types were compared from these *h* matrices using Wilcoxon’s signed rank test or Kruskal-Wallis test with *post hoc* Dunn’s test and Bonferroni correction, all implemented with the *stats* R package v4.3.1.

#### Generation of signatures and human orthologs from mouse NMF factors

To generate gene signatures from NMF factors, we first intersected the top loading genes in selected NMF factors according to scores in the *w* matrix against the most significantly upregulated genes in target cell types in relevant perturbations, taking the top 50 genes that appeared in both lists according to loading score and adjusted p-value. Thus, for M12, we compared 20-day IL-1 to control; for M18, we compared 7-day IL-1, LPS, and pIC to control; for M40, we compared LPS and pIC to control; and for M24, we compared 5FU day 8, G-CSF day 2, G-CSF day 4, LPS, and pIC to control. We mapped mouse gene symbols to human gene symbols using the *getLDS* function from the R package *biomaRt* v2.65.0, and we manually curated these results to search for any missing orthologs. A complete list of signatures and coefficients is shown in **Table S15**.

#### Gene set enrichment analysis in bulk RNA-seq samples

To test enrichment of mouse NMF factor genes or human ortholog signatures in bulk RNA-seq datasets, we used the *GSEA* function in *clusterProfiler* v4.8.3, and we generated plots using *enrichplot* v1.20.3. Published RNA-seq datasets were downloaded from the Gene Expression Omnibus (GEO) (**Table S13**). Where possible, author-generated lists of differentially expressed genes were used as input. Where this was unavailable, counts tables were downloaded and re-analyzed using *DESeq2* v1.40.2. If counts were unavailable, fastq files were downloaded, trimmed with *trim-galore*, and pseudo-aligned to the mouse mm10 transcript reference using *Salmon*, before differentially expressed genes were identified with *DESeq2*. If only normalized counts were available, *voom*, *lmFit*, and *eBayes* were used in the R package *limma* v3.58.1 to perform differential expression testing.

#### Prognostic evaluation of myelopoiesis signatures in human disease

We conducted survival analyses examining the impact and prognostic value of the EM signatures derived from our NMF analysis above in two cohorts of human AML patients. Data from 136 samples in the TCGA-LAML project^76^ and 273 BM aspirates at time of diagnosis from Beat AML^77,78^ with matched bulk RNA transcriptomic data and clinical survival data were retrieved from cBioPortal^120,121^. We normalized the NMF coefficients for genes in each signature to sum to unity. Then, each sample was given a score defined as the weighted average of the expression levels (in log(1+TPM) units) of the genes in the signature using the normalized NMF coefficients as weights. To analyze the relationship between these scores and other variables, we generated oncoplots using *oncoPrint* from *ComplexHeatmap*^122^ to visualize whether our signatures are correlated with other explanatory features such as age, sex, or mutation status, and performed two-sided Wilcoxon rank sum tests to quantify their associations. For analysis of AML prognosis, we first performed univariate Cox proportional hazards regression with covariates for overall survival in months on the Beat AML cohort to identify other potential mutational correlates of overall survival among single nucleotide polymorphisms that occurred in at least 5% of the samples using the *surv_fit* function from the *survival* R package^123,124^; p-values were adjusted for false discovery rates using the Benjamini-Hochberg procedure in *stats::p.adjust*. We obtained the Cox regression results on hSig24 alone and also for a multivariate analysis on hSig24 together with the mutational correlates that were found significant in the previous step. Age, sex, and prior history of malignancy or treatment were included as covariates in all of the Cox regression models, where available. We plotted Kaplan-Meier survival curves for the top 20% and bottom 20% for each signature respectively and conducted a log rank test for significance using the *ggsurvplot* function from *survminer*^125^. To isolate the role of hSigs from TP53 mutation status, we sub-setted the TCGA-LAML and Beat AML cohorts for TP53 wild-type (WT) patients. For these sub-cohorts, we repeated our analysis above by incorporating our hSig scores into individual Cox regression models for overall survival, derived hazard ratios and p-values, and plotted Kaplan-Meier survival curves for hSig24 for each signature in the TCGA-LAML and Beat AML sub-cohorts separately. In order to benchmark these results, we obtained ELN2022 risk classifications for patients in the two cohorts from previously published sources^126,127^ and trained additional Cox regression models using both the ELN2022 risk, clinical covariates, and each signature score on the full cohorts. We compared their performances to a baseline model trained using only the ELN2022 risk and covariates using likelihood ratio tests and ANOVA. To compare the prognostic value of our scores to that of existing risk scores, we repeated this benchmarking analysis with other gene expression-based AML prognosis scores previously described in the literature^85–88^. Moreover, we expanded our analysis and performed similar survival analyses for other non-myeloid cancers represented in TCGA, to serve as negative controls^128,129^. We also conducted survival analyses for pediatric myeloid malignancies including 1786 cases of pediatric AML from the TARGET project^130,131^. These secondary cohorts were retrieved from the cBioPortal and NCI Genomic Data Commons^132,133^. Finally, we also evaluated these myelopoiesis signature scores on bulk RNA-sequencing data from other previously reported cohorts totaling 123 adult AML patients at diagnosis without matched survival information and stratified the score distributions using a previous molecular AML classification scheme based on cell type deconvolution^75^.

#### Quantification and Statistical Analysis

Data are represented as means ± standard deviations (S.D.), or as violin plots with the center line representing the median, using R studio for molecular data or GraphPad Prism (v10.4.2) for all other data. Circles on bar graphs represent biological replicates. For experimental data, Student’s t-test was used when 2 groups were compared. Either one-way ANOVA with Tukey’s post-hoc test, or Kruskal-Wallis test with Dunn’s post-hoc test were used to compare 3 or more groups.

## Notes

https://github.com/RabadanLab/HemaScribe

### Summary of Updates

One of the author names was misspelled.

## REFERENCES

1. Olson OC, Kang YA, Passegué E (2020). Normal hematopoiesis is a balancing act of self-renewal and regeneration. Cold Spring Harb Perspect Med 10:a035519.

2. Swann JW, Olson OC, Passegué E (2024). Made to order: emergency myelopoiesis and demand-adapted innate immune cell production. Nat Rev Immunol 24:596–613.

3. Passegué E, Wagers AJ, Giuriato S, Anderson WC, Weissman IL (2005). Global analysis of proliferation and cell cycle gene expression in the regulation of stem and progenitor cell fates. J Exp Med 202:1599–1611.

4. Walter D, Lier A, Geiselhart A, Thalheimer FB, Huntscha S, Sobotta MC, Moehrle B, Brocks D, Bayindir I, Kaschutnig P, Muedder K, Klein C, Jauch A, Schroeder T, Geiger H, Dick TP, Holland-Letz T, Schmezer P, Lane SW, Rieger MA, Essers MAG, Williams DA, Trumpp A, Milsom MD (2015). Exit from dormancy provokes DNA-damage-induced attrition in haematopoietic stem cells. Nature 520:549–552.

5. Karigane D, Kobayashi H, Morikawa T, Ootomo Y, Sakai M, Nagamatsu G, Kubota Y, Goda N, Matsumoto M, Nishimura EK, Saga T, Otsu K, Suematsu M, Okamoto S, Suda T, Takubo K (2016). p38a activates purine metabolism to initiate hematopoietic stem/progenitor cell cycling in response to stress. Cell Stem Cell 19:192–204.

6. Itoh-Nakadai A, Matsumoto M, Kato H, Sasaki J, Uehara Y, Sato Y, Ebina-Shibuya R, Morooka M, Funayama R, Nakayama K, Ochiai K, Muto A, Igarashi K (2017). A Bach2-Cebp gene regulatory network for the commitment of multipotent hematopoietic progenitors. Cell Rep 18:2401–2414.

7. Kang YA, Pietras EM, Passegué E (2020). Deregulated Notch and Wnt signaling activates early-stage myeloid regeneration pathways in leukemia. J Exp Med 217:jem.20190787.

8. Pietras EM, Mirantes-Barbeito C, Fong S, Loeffler D, Kovtonyuk LV, Zhang SY, Lakshminarasimhan R, Chin CP, Techner JM, Will B, Nerlov C, Steidl U, Manz MG, Schroeder T, Passegué E (2016). Chronic interleukin-1 exposure drives haematopoietic stem cells towards precocious myeloid differentiation of the expense of self-renewal. Nat Cell Biol 18:607–618.

9. Yamashita M, Passegué E (2019). TNF-a coordinates hematopoietic stem cell survival and myeloid regeneration. Cell Stem Cell 25:357–372.

10. Reynaud D, Pietras E, Barry-Holson K, Mir A, Binnewies M, Jeanne M, Sala-Torra O, Radich JP, Passegué E (2011). IL-6 controls leukemic multipotent progenitor cell fate and contributes to chronic myelogenous leukemia development. Cancer Cell 20:661–673.

11. Welner RS, Amabile G, Bararia D, Czibere A, Yang H, Zhang H, Pontes LL, Ye M, Levantini E, Di Ruscio A, Martinelli G, Tenen DG (2015). Treatment of chronic myelogenous leukemia by blocking cytokine alterations found in normal stem and progenitor cells. Cancer Cell 27:671–81.

12. Kang YA, Paik H, Zhang SY, Chen JJ, Olson OC, Mitchell CA, Collins A, Swann JW, Warr MR, Fan R, Passegué E (2023). Secretory MPP3 reinforce myeloid differentiation trajectory and amplify myeloid cell production. J Exp Med 220:jem20230088.

13. Hérault A, Binnewies M, Leong S, Calero-Nieto FJ, Zhang SY, Kang YA, Wang X, Pietras EM, Chu SH, Barry-Holson K, Armstrong S, Gottgens B, Passegué E (2017). Myeloid progenitor cluster formation drives emergency and leukaemic myelopoiesis. Nature 544:53–58.

14. Fanti AK, Busch K, Greco A, Wang X, Cirovic B, Shang F, Nizharadze T, Frank L, Barile M, Feyerabend TB, Hofer T, Rodewald HR (2023). Flt3- and Tie2-Cre tracing identifies regeneration in sepsis from multipotent progenitors but not hematopoietic stem cells. Cell Stem Cell 30:207–218.

15. Munz CM, Dressel N, Chen M, Grinenko T, Roers A, Gerbaulet A (2023). Regeneration after blood loss and acute inflammation proceeds without contribution of primitive HSCs. Blood 141:2483–2492.

16. ImmGen Consortium (2006). Open-source ImmGen: mononuclear phagocytes. Nat Immunol 17:741.

17. Choi J, Baldwin TM, Wong M, Bolden JE, Fairfax KA, Lucas EC, Cole R, Biben C, Morgan C, Ramsay KA, Ng AP, Kauppi M, Corcoran LM, Shi W, Wilson N, Wilson MJ, Alexander WS, Hilton DJ, de Graaf CA (2019). Haemopedia RNA-seq: a database of gene expression during haematopoiesis in mice and humans. Nucleic Acids Res 47:D780–D785.

18. Zhu YP, Padgett L, Dinh HQ, Marcovecchio P, Blatchley A, Wu R, Ehinger E, Kim C, Mikilski Z, Seumois G, Madrigal A, Vijayanand P, Hedrick CC (2018) Identification of an early unipotent neutrophil progenitor with pro-tumoral activity in mouse and human bone marrow. Cell Rep 24:2329–2341.

19. Pronk CJH, Rossi DJ, Mansson R, Attema JL, Norddahl GL, Chan CKF, Sigvardsson M, Weissman IL, Bryder D (2007) Elucidation of the phenotypic, functional, and molecular topography of a myeloerythroid progenitor cell hierarchy. Cell Stem Cell 1:428–442.

20. Cabezas-Wallscheid N, Klimmeck D, Hansson J, Lipka DB, Reyes A, Wang Q, Weichenhan D, Lier A, von Paleske L, Renders S, Wunsche P, Zeisberger P, Brocks D, Gu L, Herrmann C, Haas S, Essers MAG, Brors B, Eils R, Huber W, Milsom MD, Plass C, Krijgsveld J, Trumpp A (2014) Identification of regulatory networks in HSCs and their immediate progeny via integrated proteome, transcriptome, and DNA methylome analysis. Cell Stem Cell 15:507–522.

21. Pietras EM, Reynaud D, Kang YA, Carlin D, Calero-Nieto FJ, Leavitt AD, Stuart JM, Gottgens B, Passegué E (2015). Functionally distinct subsets of lineage-biased multipotent progenitors control blood production in normal and regenerative conditions. Cell Stem Cell 17:35–46.

22. Yanez A, Coetzee SG, Olsson A, Muench DE, Berman BP, Hazelett DJ, Salomonis N, Grimes HL, Goodridge HS (2017). Granulocyte-monocyte progenitors and monocyte-dendritic cell progenitors independently produce functionally distinct monocytes. Immunity 47:890–902.

23. Collins A, Swann JW, Proven MA, Patel CM, Mitchell CA, Kasbekar M, Dellorusso PV, Passegué E (2024). Maternal inflammation regulates fetal emergency myelopoiesis. Cell 187:1402–1421.

24. Klein F, Roux J, Cvijetic G, Fernandes Rodrigues P, von Muenchow L, Lubin R, Pelczar P, Yona S, Tsapogas P, Tussiwand R (2022). Dntt expression reveals developmental hierarchy and lineage specification of hematopoietic progenitors. Nat Immunol 23:505–517.

25. Izzo F, Lee SC, Poran A, Chaligne R, Gaiti F, Gross B, Murali RR, Deochand SD, Ang C, Jones PW, Nam AS, Kim KT, Kothen-Hill S, Schulman RC, Ki M, Lhoumaud P, Skok JA, Viny AD, Levine RL, Kenigsberg E, Abdel-Wahab O, Landau DA (2020). DNA methylation disruption reshapes the hematopoietic differentiation landscape. Nat Genet 52:378–387.

26. Tabula Muris Consortium (2018). Single-cell transcriptomics of 20 mouse organs creates a Tabula Muris. Nature 562:367–372.

27. Swann JW, Zhang R, Verovskaya EV, Calero-Nieto FJ, Wang X, Proven MA, Shyu PT, Guo XE, Gottgens B, Passegué E (2024). Inflammation perturbs hematopoiesis by remodeling specific compartments of the bone marrow niche. bioRxiv doi:10.1101/2024.09.12.612751.

28. Carreira-Perpiñán MA (2010). The elastic embedding algorithm for dimensionality reduction. 27^th^ International Conference on Machine Learning, Haifa. 10:167–174.

29. Chen Z, An S, Bai X, Gong F, Ma L, Wan L (2019). DensityPath: an algorithm to visualize and reconstruct cell state-transition path on density landscape for single-cell RNA sequencing data. Bioinformatics 35:2593–2601.

30. Wolf FA, Hamey FK, Plass M, Solana J, Dahlin JS, Gottgens B, Rajewsky N, Simon L, Theis FJ (2019). PAGA: graph abstraction reconciles clustering with trajectory inference through a topology preserving map of single cells. Genome Biol 20:59

31. Kucinski I, Campos J, Barile M, Severi F, Bohin N, Moreira PN, Allen L, Lawson H, Haltalli MLR, Kinston SJ, O’Carroll D, Kranc KR, Gottgens B (2024). A time- and single-cell-resolved model of murine bone marrow hematopoiesis. Cell Stem Cell 31:244–259.

32. Aibar S, Gonzalez-Blas CB, Moerman T, Huynh-Thu VA, Imrichova H, Hulselmans G, Rambow F, Marine JC, Geurts P, Aerts J, van den Oord J, Atak ZK, Wouters J, Aerts S (2017). SCENIC: single-cell regulatory network inference and clustering. Nat Methods 14:1083–1086.

33. Pevny L, Simon MC, Robertson E, Klein WH, Tsai SF, D’Agati V, Orkin SH, Costantini F (1991). Erythroid differentiation in chimaeric mice blocked by a targeted mutation in the gene for transcription factor GATA-1. Nature 349:257–260.

34. Zhang DE, Zhang P, Wang ND, Hetherington CJ, Darlington GJ, Tenen DG (1997). Absence of granulocyte colony-stimulating factor signaling and neutrophil development in CCAAT enhancer binding protein alpha-deficient mice. PNAS 94:569–574.

35. Bergqvist I, Eriksson M, Saarikettu J, Eriksson B, Corneliussen B, Grundstrom T, Holmberg D (2000). The basic helix-loop-helix transcription factor E2-2 is involved in T lymphocyte development. Eur J Immunol 30:2857–2863.

36. Bertero A, Vallier L (2015). Fucci2a mouse upgrades live cell cycle imaging. Cell Cycle 14:948–949.

37. Setty M, Kiseliovas V, Levine J, Gayoso A, Mazutis L, Pe’er D (2019). Characterization of cell fate probabilities in single-cell data with Palantir. Nat Biotechnol 37:451–460.

38. Rodriguez-Fraticelli A, Wolock SL, Weinreb CS, Panero R, Patel SH, Jankovic M, Sun J, Calogero RA, Klein AM, Camargo FD (2018). Clonal analysis of lineage fate in native haematopoiesis. Nature 553:212–216.

39. Dellorusso PV, Proven MA, Calero-Nieto FJ, Wang X, Mitchell CA, Hartmann F, Amouzgar M, Favaro P, DeVilbiss A, Swann JW, Ho TT, Zhao Z, Bendall SC, Morrison S, Gottgens B, Passegué E (2024). Autophagy counters inflammation-driven glycolytic impairment in aging hematopoietic stem cells. Cell Stem Cell 31:1020–1037.

40. Pietras EM, Lakshminarasimhan R, Techner JM, Fong S, Flach J, Binnewies M, Passegué E (2014). Re-entry into quiescence protects hematopoietic stem cells from the killing effect of chronic exposure to type I interferons. J Exp Med 211:245–262.

41. Tusi BK, Wolock SL, Weinreb CS, Hwang Y, Hidalgo D, Zilionis R, Waisman A, Huh JR, Klein AM, Socolovsky M (2018). Population snapshots predict early haematopoietic and erythroid hierarchies. Nature 555:54–60.

42. Mitchell CA, Verovskaya EV, Calero-Nieto FJ, Olson OC, Swann JW, Wang X, Herault A, Dellorusso PV, Zhang SY, Svendsen AF, Pietras EM, Bakker ST, Ho TT, Gottgens B, Passegué E (2023). Stromal niche inflammation mediated by IL-1 signalling is a targetable driver of haematopoietic ageing. Nat Cell Biol 25:30–41.

43. Cao Y, Lin Y, Ormerod JT, Yang P, Yang JYH, Lo KK (2019). scDC: single cell differential composition analysis. BMC Bioinformatics 20:721.

44. Skinnider MA, Squair JW, Kathe C, Anderson MA, Gautier M, Matson KJE, Milano M, Hutson TH, Barraud Q, Philips AA, Foster LJ, La Manno G, Levine AJ, Courtine G (2021). Cell type prioritization in single-cell data. Nat Biotechnol 39:30–34.

45. Li JJ, Liu J, Li YE, Chen LV, Cheng H, Li Y, Cheng T, Wang QF, Zhou BO (2024). Differentiation route determines the functional outputs of adult megakaryopoiesis. Immunity 57:478–494.

46. Poscablo DM, Worthington AK, Smith-Berdan S, Rommel MGE, Manso BA, Adili R, Mok L, Reggiardo RE, Cool T, Mogharrab R, Myers J, Dahmen S, Medina P, Beaudin AE, Boyer SW, Holinstat M, Jonsson VD, Forsberg EC (2024). An age-progressive platelet differentiation path from hematopoietic stem cells causes exacerbated thrombosis. Cell 187:3090–3107.

47. Haas S, Hansson J, Klimmeck D, Loeffler D, Velten L, Uckelmann H, Wurzer S, Prendergast AM, Schnell A, Hexel K, Santarella-Mellwig R, Blasxkiewicz S, Kuck A, Geiger H, Milsom MD, Steinmetz LM, Schroeder T, Trumpp A, Krijgsveld J, Essers MAG (2015). Inflammation-induced emergency megakaryopoiesis driven by hematopoietic stem cell-like megakaryocyte progenitors. Cell Stem Cell 17:422–434.

48. Carrelha J, Mazzi S, Winroth A, Hagemann-Jensen M, Ziegenhain K, Seki M, Brennan MS, Lehander M, Wu B, Meng Y, Markljung E, Norfo R, Ishida H, Stralin KB, Grasso F, Karali CS, Aliouat A, Hillen A, Chari E, Siletti K, Thongjuea S, Mead AJ, Linarsson S, Nerlov C, Sandberg R, Yoshizato T, Woll PS, Jacobsen SEW (2024). Alternative platelet differentiation pathways initiated by nonhierarchically related hematopoietic stem cells. Nat Immunol 25:1007–1019.

49. Wright DE, Cheshier SH, Wagers AJ, Randall TD, Christensen JL, Weissman IL (2001). Cyclophosphamide/granulocyte colony-stimulating factor causes selective mobilization of bone marrow hematopoietic stem cells into the blood after M phase of the cell cycle. Blood 97:2278–2285.

50. Athanasiou M, Mavrothalassitis G, Sun-Hoffman L, Blair DG (2000). FLI-1 is a suppressor of erythroid differentiation in human hematopoietic cells. Leukemia 14:439–445.

51. Balogh P, Adelman ER, Pluvinage JV, Capaldo BJ, Freeman KC, Singh S, Elagib KE, Nakamura Y, Kurita R, Sashida G, Zunder ER, Li H, Gru AA, Price EA, Schrier SL, Weissman IL, Figueroa ME, Pang WW, Goldfarb AN (2020). RUNX3 levels in human hematopoietic progenitors are regulated by aging and dictate erythroid-myeloid balance. Haematologica 105:905–913.

52. Growney JD, Shigematsu H, Li Z, Lee BH, Adelsperger J, Rowan R, Curley DP, Kutok JL, Akashi K, Williams IR, Speck NA, Gilliland DG (2005). Loss of Runx1 perturbs adult hematopoiesis and is associated with a myeloproliferative phenotype. Blood 106:494–504.

53. Huang G, Zhang P, Hirai H, Elf S, Yan X, Chen Z, Koschmieder S, Okuno Y, Dayaram T, Growney JD, Shivdasani RA, Gilliland DG, Speck NA, Nimer SD, Tenen DG (2008). PU.1 is a major downstream target of AML1 (RUNX1) in adult mouse hematopoiesis. Nat Genet 40:51–60.

54. Zhao X, Bartholdy B, Yamamoto Y, Evans EK, Alberich-Jorda M, Staber PB, Benoukraf T, Zhang P, Zhang J, Trinh BQ, Crispino JD, Hoang T, Bassal MA, Tenen DG (2022). PU.1-c-Jun interaction is crucial for PU.1 function in myeloid development. Commun Biol 14:961.

55. Cui A, Huang T, Li S, Ma A, Perez JL, Sander C, Keskin DB, Wu CJ, Fraenkel E, Hacohen N (2024). Dictionary of immune responses to cytokines at single-cell resolution. Nature 625:377–384.

56. Bouman BJ, Demerdash Y, Sood S, Grunschlager F, Pilz F, Itani AR, Kuck A, Marot-Lassauzaie V, Haas S, Haghverdi L, Essers MA (2023). Single-cell time series analysis reveals the dynamics of HSPC response to inflammation. Life Sci Alliance 7:e202302309.

57. Fast EM, Sporrij A, Manning M, Rocha EL, Yang S, Zhou Y, Guo J, Baryawno N, Barkas N, Scadden D, Camargo F, Zon LI (2021). External signals regulate continuous transcriptional states in hematopoietic stem cells. Elife 10:e66512.

58. McClatchy J, Strogantsev R, Wolfe E, Lin HY, Mohammadhosseini M, Davis BA, Eden C, Goldman D, Fleming WH, Conley P, Wu G, Cimmino L, Mohammed H, Agarwal A (2023). Clonal hematopoiesis related Tet2 loss-of-function impedes IL-1β-mediated epigenetic reprogramming in hematopoietic stem and progenitor cells. Nat Commun 14:8102.

59. Zhang S, Paccalet A, Rohde D, Cremer S, Hulsmans M, Lee IH, Mentkowski K, Grune J, Schloss MJ, Honold L, Iwamoto Y, Zheng Y, Bredella MA, Buckless C, Ghoshhajra B, Thondapu V, van der Laan AM, Piek JJ, Niessen HWM, Pallante F, Carnevale R, Perrotta S, Carnevale D, Iborra-Egea O, Munoz-Guijosa C, Galvez-Monton C, Bayes-Genis A, Vidoudez C, Trauger SA, Scadden D, Swirski FK, Moskowitz MA, Naxerova K, Nahrendorf M (2023). Bone marrow adipocytes fuel emergency hematopoiesis after myocardial infarction. Nat Cardiovasc Res 2:1277–1290.

60. Robles-Vera I, Jarit-Cabanillas A, Brandi P, Martinez-Lopez M, Martinez-Cano S, Rodrigo-Tapias M, Femenia-Muina M, Redondo-Urzainqui A, Nunez V, Gonzalez-Correa C, Moleon J, Duarte J, Conejero L, Mata-Martinez P, Diez-Rivero CM, Bergon-Gutierrez M, Fernandez-Lopez I, Gomez MJ, Quintas A, Dopazo A, Sanchez-Cabo F, Pariente E, Del Fresno C, Subiza JL, Iborra S, Sancho D (2025). Microbiota translocation following intestinal barrier disruption promotes Mincle-mediated training of myeloid progenitor in the bone marrow. Immunity 58:381–396.

61. Regan-Komito D, Swann JW, Demetriou P, Cohen ES, Horwood NJ, Sansom SN, Griseri T (2020). GM-CSF drives dysregulated hematopoietic stem cell activity and pathogenic extramedullary myelopoiesis in experimental spondyloarthritis. Nat Commun 11:155.

62. Ikeda N, Kubota H, Suzuki R, Morita M, Yoshimura A, Osada Y, Kishida K, Kitamura D, Iwata A, Yotsumoto S, Kurotaki D, Nishimura K, Nishiyama A, Tamura T, Kamatani T, Tsunoda T, Murakawa M, Asahina Y, Hayashi Y, Harada H, Harada Y, Yokota A, Hirai H, Seki T, Kuwahara M, Yamashita M, Shichino S, Tanaka M, Asano K (2023). The early neutrophil-committed progenitors aberrantly differentiate into immunoregulatory monocytes during emergency myelopoiesis. Cell Rep 42:112165.

63. Rodriguez-Fraticelli A, Weinreb C, Wang SW, Migueles RP, Jankovic M, Usart M, Klein AM, Lowell S, Camargo FD (2020). Single-cell lineage tracing unveils a role for TCF15 in haematopoiesis. Nature 583:585–589.

64. Kamimoto K, Stringa B, Hoffmann CM, Jindal K, Solnica-Krezel L, Morris SA (2023). Dissecting cell identity via network inference and in silico gene perturbation. Nature 614:742–751.

65. Alkrekshi A, Wang W, Rana PS, Markovic V, Sossey-Alaoui K (2021). A comprehensive review of the functions of YB-1 in cancer stemness, metastasis and drug resistance. Cell Signal 85:110073.

66. Tailor D, Resendez A, Garcia-Marques FJ, Pandrala M, Going CC, Bermudez A, Kumar V, Rafat M, Nambiar DK, Honkala A, Le QT, Sledge GW, Graves E, Pitteri SJ, Malhotra SV (2021). Y box binding protein 1 inhibition as a targeted therapy for ovarian cancer. Cell Chem Biol 28:1206–1220.

67. Antony-Debré I, Paul A, Leite J, Mitchell K, Kim HM, Carvajal LA, Todorova TI, Huang K, Kumar A, Farahat AA, Bartholdy B, Narayanagari SR, Chen J, Ambesi-Impiombato A, Ferrando AA, Mantzaris I, Gavathiotis E, Verma A, Will B, Boykin DW, Wilson WD, Poon GM, Steidl U (2017). Pharmacological inhibition of the transcription factor PU.1 in leukemia. J Clin Invest 127:4297–4313.

68. You G, Zhang M, Bian Z, Guo H, Xu Z, Ni Y, Lan Y, Yue W, Gong Y, Chang Y, Huang X, Lui B (2022). Decoding lymphomyeloid divergence and immune hyporesponsiveness in G-CSF-primed human bone marrow by single-cell RNA-seq. Cell Discov 8:59.

69. Reyes M, Filbin MR, Bhattacharyya RP, Billman K, Eisenhaure T, Hung DT, Levy BD, Baron RM, Blainey PC, Goldberg MB, Hacohen N (2020). An immune-cell signature of bacterial sepsis. Nat Med 26:333–340.

70. Zeng AGX, Iacobucci I, Shah S, Mitchell A, Wong G, Bansal S, Chen D, Gao Q, Kim H, Kennedy JA, Arruda A, Minden MD, Haferlach T, Mullighan CG, Dick JE (2025) Single-cell Transcriptional atlas of human hematopoiesis reveals genetic and hierarchy-based determinants of aberrant AML differentiation. Blood Cancer Discov. OF1–OF18.

71. Wilk AJ, Lee MJ, Wei B, Parks B, Pi R, Martinez-Colon GJ, Ranganath T, Zhao NQ, Taylor S, Becker W, Stanford COVID-19 Biobank, Jimenez-Morales D, Blomkalns AL, O’Hara R, Ashley EA, Nadeau KC, Yang S, Holmes S, Rabinovitch M, Rogers AJ, Greenleaf WJ, Blish CA (2021). Multi-omic profiling reveals widespread dysregulation of innate immunity and hematopoiesis in COVID-19. J Exp Med 218:e20210582.

72. Boroujeni ME, Sekrecka A, Antonczyk A, Hassani S, Sekrecki M, Nowicka H, Lopacinska N, Olya A, Kluzek K, Wesoly J, Bluyssen HAR (2022). Dysregulated interferon response and immune hyperactivation in severe COVID-19: targeting STATs as a novel therapeutic strategy. Front Immunol 13:888897.

73. Krivtsov AV, Figueroa ME, Sinha AU, Stubbs MC, Feng Z, Valk PJM, Delwel R, Dohner K, Bullinger L, Kung AL, Melnick AM, Armstrong SA (2013). Cell of origin determines clinically relevant subtypes of MLL-rearranged AML. Leukemia 27:852–860.

74. Shih AH, Jiang Y, Meydan C, Shank K, Pandey S, Barreyro L, Antony-Debre I, Viale A, Socci N, Sun Y, Robertson A, Cavatore M, de Stanchina E, Hricik T, Rapaport F, Woods B, Wei C, Hatlen M, Baljevic M, Nimer SD, Tallman M, Paietta E, Cimmino L, Aifantis I, Steidl U, Mason C, Melnick A, Levine RL (2015). Mutational cooperativity linked to combinatorial epigenetic gain of function in acute myeloid leukemia. Cancer Cell 27:502–515.

75. Zeng AGX, Bansal S, Jin L, Mitchell A, Chen WC, Abbas HA, Chan-Seng-Yue M, Voisin V, van Galen P, Tierens A, Cheok M, Preudhomme C, Dombret H, Daver N, Futreal PA, Minden MD, Kennedy JA, Wang JCY, Dick JE (2022). A cellular hierarchy framework for understanding heterogeneity and predicting drug response in acute myeloid leukemia. Nat Med 28:1212–1223.

76. The Cancer Genome Atlas Research Network, et al. (2013). Genomic and epigenomic landscapes of adult de novo acute myeloid leukemia. N Engl J Med 368:2059–2074.

77. Tyner JW, Tognon CE, Bottomly D, et al. (2018). Functional genomic landscape of acute myeloid leukaemia. Nature 562:526–531.

78. Bottomly D, Long N, Schultz AR, et al. (2022). Integrative analysis of drug response and clinical outcome in acute myeloid leukemia. Cancer Cell 40:850–864.

79. Papaemmanuil E, Gerstung M, Bullinger L, Gaidzik VI, Paschka P, Roberts ND, Potter NE, Heuser M, Thol F, Bolli N, Gundem G, Van Loo P, Martincorena I, Ganly P, Mudie L, McLaren S, O’Meara S, Raine K, Jones DR, Teague JW, Butler AP, Greaves MF, Ganser A, Döhner K, Schlenk RF, Döhner H, Campbell PJ (2016). Genomic classification and prognosis in acute myeloid leukemia. N Engl J Med 374:2209–2221.

80. Aung MMK, Mills ML, Bittencourt-Silvestre J, Keeshan K (2021) Insights into the molecular profiles of adult and paediatric acute myeloid leukaemia. Mol Oncol 15:2253–2272.

81. The Cancer Genome Atlas Research Network, et al. (2014). Comprehensive molecular profiling of lung adenocarcinoma. Nature 511:543–550.

82. Zhu Q, Chai Y, Jin L, Ma Y, Lu H, Chen Y, Feng W (2023). Construction and validation of a novel prognostic model of neutrophil-related genes signature of lung adenocarcinoma. Sci Rep 13:18226.

83. Wu D, Liu Y, Liu J, Ma L, Tong X (2024). Myeloid cell differentiation-related gene signature for predicting clinical outcome, immune microenvironment, and treatment response in lung adenocarcinoma. Sci Rep 14:17460.

84. Dohner H, Wei AH, Appelbaum FR, Craddock C, DiNardo CD, Dombret H, Ebert BL, Fenaux P, Godley LA, Hasserjian RP, Larson RA, Levine RL, Miyazaki Y, Niederwieser D, Ossenkoppele G, Rollig C, Sierra J, Stein EM, Tallman MS, Tien HF, Wang J, Wierzbowska A, Lowenberg B (2022). Diagnosis and management of AML in adults: 2022 recommendations from an international expert panel on behalf of the ELN. Blood 140:1345–1377.

85. Nehme A, Dakik H, Picou F, Cheok M, Preudhomme C, Dombret H, Lambert J, Gyan E, Pigneux A, Recher C, Bene MC, Gouilleux F, Zibara K, Herault O, Mazurier F (2020). Horizontal meta-analysis identifies common deregulated genes across AML subgroups providing a robust prognostic signature. Blood Adv 4:5322–5335.

86. Docking TR, Parker JDK, Jadersten M, Duns G, Chang L, Jiang J, Pilsworth JA, Swanson LA, Chan SK, Chiu R, Nip KM, Mar S, Mo A, Wang X, Martinez-Hoyer S, Stubbins RJ, Mungall KL, Mungall AJ, Moore RA, Jones SJM, Birol I, Marra MA, Hogge D, Karsan A (2021). A clinical transcriptome approach to patient stratification and therapy selection in acute myeloid leukemia. Nat Commun 12:2474.

87. Ng SWK, Mitchell A, Kennedy JA, Chen WC, McLeod J, Ibrahimova N, Arruda A, Popescu A, Gupta V, Schimmer AD, Schuh AC, Yee KW, Bullinger L, Herold T, Gorloch D, Buchner T, Hiddemann W, Berdel WE, Wormann B, Cheok M, Preudhomme C, Bombret H, Metzeler K, Buske C, Lowenberg B, Valk PJM, Zandstra PW, Minden MD, Dick JE, Wang JCY (2016). A 17-gene stemness score for rapid determination of risk in acute leukaemia. Nature 540:433–437.

88. Isobe T, Kuciniski I, Barile M, Wang X, Hannah R, Bastos HP, Chabra S, Vijayabaskar MS, Sturgess KHM, Williams MJ, Giotopoulos G, Marando L, Li J, Rak J, Gozdecka M, Prins D, Shepherd MS, Watcham S, Green AR, Kent DG, Vassiliou GS, Huntly BJP, Wilson NK, Gottgens B (2023). Preleukemic single-cell landscapes reveal mutation-specific mechanisms and gene programs predictive of AML patient outcomes. Cell Genom 3:100426.

89. Laurenti E, Gottgens B (2018) From haematopoietic stem cells to complex differentiation landscapes. Nature 553:418–426.

90. Drissen R, Buza-Vidas N, Woll P, Thongjuea S, Gambardella A, Giustacchini A, Mancini E, Zriwil A, Lutteropp M, Grover A, Mead A, Sitnicka E, Jacobsen SEW, Nerlov C (2016). Distinct myeloid progenitor-differentiation pathways identified through single-cell RNA sequencing. Nat Immunol 17:666–676.

91. Kwok AJ, Allcock A, Ferreria RC, Cano-Gamez E, Smee M, Burnham KL, Zurke YX, Emergency Medicine Research Oxford (EMROx), McKechnie S, Mentzer AL, Monaco C, Udalova IA, Hinds CJ, Todd JA, Davenport EE, Knight JC (2023). Neutrophils and emergency granulopoiesis drive immune suppression and an extreme response endotype drugin sepsis. Nat Immunol 24:767–779.

92. Scheiermann C, Frenette PS, Hidalgo A (2015). Regulation of leucocyte homeostasis in the circulation. Cardiovasc Res 107:340–351.

93. Shultz AJ, Sackton TB (2019). Immune genes are hotspots of shared positive selection across birds and mammals. Elife 8:e41815.

94. Lin KK, Rossi L, Boles NC, Hall BE, George TC, Goodell MA (2011). CD81 is essential for re-entry of hematopoietic stem cells to quiescence following stress-induced proliferation via deactivation of the Akt pathway. PLoS Biol 9:e1001148.

95. Ichikawa M, Yoshimi A, Nakagawa M, Nishimoto N, Watanabe-Okochi N, Kurokawa M (2013). A role for RUNX1 in hematopoiesis and myeloid leukemia. Int J Hematol 97:726–734.

96. Larsen SB, Cowley CJ, Sajjath SM, Barrows D, Yang Y, Carroll TS, Fuchs E (2021) Establishment, maintenance, and recall of inflammatory memory. Cell Stem Cell 28:1758–1774.

97. Perner F, Schnoeder TM, Xiong Y, Jayavelu AK, Mashamba N, Santamaria NT, Huber N, Todorova K, Hatton C, Perner B, Eifert T, Murphy C, Hartmann M, Hoell JI, Schroder N, Brandt S, Hochhaus A, Mertens PR, Mann M, Armstrong SA, Mandinova A, Heidel FH (2022). YBX1 mediates translation of oncogenic transcripts to control cell competition in AML. Leukemia 36:426–437.

98. Schnoeder TM, Perner F, Jayavelu AK, Mao L, Zhang Q, Hsu CJ, Eifert T, Grunwald U, Mertens P, Tailor D, Taherinasab A, Braun TP, Malhotra S, Heidel FH (2022). Pre-clinical investigation of a novel small molecule inhibitor targeting Ybx1 in AML. Blood 140:491–492.

99. Bommert KS, Effenberger M, Leich E, Kuspert M, Murphy D, Langer C, Moll R, Janz S, Mottok A, Weissbach S, Rosenwald A, Bargou R, Bommert K (2013). The feed-forward loop between YB-1 and MYC is essential for multiple myeloma cell survival. Leukemia 27:441–450.

## Additional references for methods

100. Heumos L, Schaar AC, Lance C, Litinetskaya A, Drost F, Zappia L, Lucken MD, Strobl DC, Henao J, Curion F, Single-cell best practices consortium, Schiller HB, Theis FJ (2023). Best practices for single-cell analysis across modalities. Nat Rev Genet 24:550–572.

101. McGinnis CS, Patterson DM, Winkler J, Conrad DN, Hein MY, Srivastava V, Hu JL, Murrow LM, Weissman JS, Werb Z, Chow ED, Gartner ZJ (2019). MULTI-seq: sample multiplexing for single-cell RNA sequencing using lipid-tagged indices. Nat Methods 16:619–626.

102. Hao Y, Hao S, Andersen-Nissen E, Mauck WM, Zhang S, Butler A, Lee MJ, Wilk AJ, Darby C, Zager M, Hoffman P, Stoeckius M, Papalexi E, Mimitou EP, Jain J, Srivastava A, Stuart T, Fleming LM, Yeung B, Rogers AJ, McElrath JM, Blish CA, Gottardo R, Smibert P, Satija R (2021). Integrated analysis of multimodal single-cell data. Cell 184:3573–3587.

103. Hao Y, Stuart T, Kowalski MH, Choudhary S, Hoffman P, Hartman A, Srivastava A, Molla G, Madad S, Fernandez-Granda C, Satija R (2024). Dictionary learning for integrative, multimodal and scalable single-cell analysis. Nat Biotechnol 42:293–304.

104. Germain PL, Lun A, Mexide CG, Macnair W, Robinson MD (2021) Doublet identification in single-cell sequencing data using scDblFinder. F1000Res 10:979.

105. Andreatta M, Carmona SJ (2021). UCell: robust and scalable single-cell gene signature scoring. Comput Struct Biotechnol 19:3796–3798.

106. Aran D, Looney AP, Liu L, Wu E, Fong V, Hsu A, Chak S, Nakiwadi RP, Wolters PJ, Abate AR, Butte AJ, Bhattacharya M (2019). Reference-based analysis of lung single-cell sequencing reveals a transitional profibrotic macrophage. Nat Immunol 20:163–172.

107. Stuart T, Butler A, Hoffman P, Hafmeister C, Papalexi E, Mauck WM, Hao Y, Stoeckius M, Smibert P, Satija R (2019). Comprehensive integration of single-cell data. Cell 177:1888–1902.

108. Tang M, Kaymaz Y, Logeman BL, Eichhorn S, Liang ZS, Dulac C, Sackton TB (2021). Evaluating single-cell cluster stability using the Jaccard similarity index. Bioinformatics 37:2212–2214.

109. Wasserman L (2018). Topological data analysis. Annu Rev Sta Appl 5:501–532.

110. Cheng Y (1995). Mean shift, mode seeking, and clustering. IEEE Trans Pattern Anal Mach Intell 17:790–799.

111. Comaniciu D, Meer P (2002). Mean shift: a robust approach toward feature space analysis. IEEE Trans Pattern Anal Mach Intell 24:603–619.

112. Kang JB, Nathan A, Weinand K, Zhang F, Willard N, Rumker L, Moody DB, Korsunsky I, Raychaudhuri S (2021). Efficient and precise single-cell reference atlas mapping with Symphony. Nat Commun 12:5890.

113. Pedregosa F, Varoquaux G, Gramfort A, Michel V, Thirion B, Grisel O, Blondel M, Prettenhofer P, Weiss R, Dubourg V, Vanderplas J, Passos A, Cournapeau D, Bricher M, Perrot M, Duchesnay E (2011). Scikit-learn: machine learning in Python. J Mach Learn Res 12:2825–2830.

114. Van de Sande B, Flerin C, Davie K, De Waegeneer M, Hulselmans G, Aibar S, Seurinck R, Saelens W, Cannoodt R, Rouchon Q, Verbeiren T, De Maeyer D, Reumers J, Saeys Y, Aerts S (2020). A scalable SCENIC workflow for single-cell gene regulatory network analysis. Nat Protoc 15:2247–2276.

115. Paul F, Arkin Y, Giladi A, Jaitin DA, Kenigsberg E, Keren-Shaul H, Winter D, Lara-Astiaso D, Gury M, Weiner A, David E, Cohen N, Lauridsen FKB, Haas S, Schlitzer A, Mildner A, Ginhoux F, Jung S, Trumpp A, Porse BT, Tanay A, Amit I (2015). Transcriptional heterogeneity and lineage commitment in myeloid progenitors. Cell 163:1663–1677.

116. Stuart T, Srivastava A, Madad S, Laraeu CA, Satija R (2021). Single-cell chromatin state analysis with Signac. Nat Methods 18:1333–1341.

117. DeBruine ZJ, Pospisilik JA, Triche TJ (2024). Fast and interpretable non-negative matrix factorization for atlas-scale single cell data. bioRxiv doi:10.1101/2021.09.01.458620.

118. Yu G, Wang LG, Han Y, He QY (2012). clusterProfiler: an R package for comparing biological themes among gene clusters. OMICS 16:284–287.

119. Wu T, Hu E, Xu S, Chen M, Guo P, Dai Z, Feng T, Zhou L, Tang W, Zhan L, Fu X, Liu S, Bo X, Yu G (2021). clusterProfiler 4.0: a universal enrichment tool for interpreting omics data. Innovation 2:100141.

120. Cerami E, Gao J, Dogrusoz U, Gross BE, Sumer SO, Aksoy BA, Jacobsen A, Byrne CJ, Heuer ML, Larsson E, Antipin Y, Reva B, Goldberg AP, Sander C, Schultz N (2012). The cBio Cancer Genomics Portal: an open platform for exploring multidimensional cancer genomics data. Cancer Discov 2:401–404.

121. Gao J, Aksoy BA, Dogrusoz U, Dresdner G, Gross B, Sumer SO, Sun Y, Jacobsen A, Sinha R, Larsson E, Cerami E, Sander C, Schultz N (2013). Integrative analysis of complex cancer genomic clinical profiles using the cBioPortal. Sci Signal 6:pl1.

122. Gu Z, Eils R, Schlesner M (2016). Complex heatmaps reveal patterns and correlations in multidimensional genomic data. Bioinformatics 32:2847–2849.

123. Therneau T (2024). A package for survival analysis in R. R package version 3.7–0, https://CRAN.R-project.org/package=survival.

124. Therneau TM, Grambsch PM (2000). Modeling Survival Data: Extending the Cox Model. Springer, New York. ISBN 0-387-98784-3.

125. Kassambara A, Kosinski M, Biecek P (2024). survminer: Drawing Survival Curves using ‘ggplot2’. R package version 0.5.0, https://rpkgs.datanovia.com/survminer/index.html.

126. Lachowiez CA, Long N, Saultz J, Gandhi A, Newell LF, Hayes-Lattin B, Maziarz RT, Leonard J, Bottomly D, McWeeney S, Dunlap J, Press R, Meyers G, Swords R, Cook RJ, Tyner JW, Druker BJ, Traer E (2023). Comparison and validation of the 2022 European LeukemiaNet guidelines in acute myeloid leukemia. Blood Adv 7:1899–1909.

127. Lee Y, Baughn LB, Myers CL, Sachs Z (2024). Machine learning analysis of gene expression reveals TP53 mutant-like AML with wild type TP53 and poor prognosis. Blood Cancer J 14:80.

128. The Cancer Genome Atlas Network (2012). Comprehensive molecular portraits of human breast tumours. Nature 490:61–70.

129. Ciriello G, Gatza ML, Beck AH, Wilkerson MD, Rhie SK, Pastore A, Zhang H, McLellan M, Yau C, Kandoth C, Bowlby R, Shen H, Hayat S, Fieldhouse R, Lester SC, Tse GMK, Factor RE, Collins LC, Allison KH, Chen YY, Jensen K, Johnson NB, Oesterreich S, Mills GB, Cherniak AD, Robertson G, Benz C, Sander C, Laird PW, Hoadley KA, King TA, TCGA Research Network, Perou CM (2015). Comprehensive molecular portraits of invasive lobular breast cancer. Cell 163:506–519.

130. McNeer NA, Philip J, Geiger H, Ries RE, Lavallee VP, Walsh M, Shah M, Arora K, Emde AK, Robine N, Alonzo TA, Kolb EA, Gamis AS, Smith M, Gerhard DS, Guidry-Auvil J, Meshinchi S, Kentsis A (2019). Genetic mechanisms of primary chemotherapy resistance in pediatric acute myeloid leukemia. Leukemia 33:1934–1943.

131. Bolouri H, Farrer JE, Triche T, Ries RE, Lim EL, Alonzo TA, Ma Y, Moore R, Mungall AJ, Marra MA, Zhang J, Ma X, Liu Y, Auvil JMG, Davidsen TM, Gesuwan P, Hermida LC, Salhia B, Capone S, Ramsingh G, Zwaan CM, Noort S, Piccolo SR, Kolb EA, Gamis AS, Smith MA, Gerhard DS, Meshinchi S (2019). The molecular landscape of pediatric acute myeloid leukemia reveals recurrent structural alterations and age-specific mutational interactions. Nat Med 25:530.

132. Heath AP, Ferretti V, Agrawal S, et al (2021). The NCI Genomics Data Commons. Nat Genet 53:257–262.

133. De Bruijn I, Kundra R, Mastrogiacomo B, et al (2023). Analysis and visualization of longitudinal genomic and clinical data from the AACR project GENIE Biopharma Collaborative in cBioPortal. Cancer Res 83:3861–3867.

